# A Cambrian soft-bodied conical animal illustrates the origins of lophophorate phyla

**DOI:** 10.1101/2022.03.19.485005

**Authors:** Han Zeng, Xiangyuan Chen, Yao Liu, Maoyan Zhu, Fangchen Zhao, Aihua Yang

## Abstract

The origin and early evolution of lophotrochozoans remain a difficult but crucial issue in reconstructing metazoan phylogeny. Exceptionally preserved fossils have provided hopeful information for resolving this lophotrochozoan problem. Here we identify that *Conicula striata*, a soft-bodied conical animal from the early Cambrian Chengjiang Lagerstätte, has mosaic characteristics of different lophophorate phyla. *C. striata* possesses a phoronid-like vermiform trunk housing a U-shaped gut and a coiled lophophore comprising ≥12 arms with numerous tentacles, but it also bears brachiopod-like features including an undivided mantle enclosing lophophore and a pedicle-like protrusion on annulated trunk. Phylogenetic analysis retrieves *C. striata* as a total-group lophophorate and as an intermediate taxon between Phoronida and Brachiopoda. This suggests that the bivalved architecture of brachiopods originated from an undivided mantle in a phoronid-like ancestor, and that the ancestral vermiform trunk became reduced during the origin of Brachiopoda, illuminating the origins of body plans in the lophophorate phyla.

## Introduction

The lophophorates, which comprise three extant metazoan phyla including Brachiopoda, Phoronida, and Bryozoa (Ectoprocta), are a major group of lophotrochozoans featuring a tentacular filter-feeding apparatus called lophophore (Hyman, 1959; Brusca *et al*., 2016). The origins of lophophorate phyla have attracted interest from evolutionary biologists for decades but remain unresolved (Emig, 1984; Valentine, 2004; Dunn *et al*., 2014). The lophophorates have long been considered as monophyletic (Lophophorata) based on their shared body plans (Caldwell, 1882; Hatschek, 1888; Hyman, 1959; Emig, 1984). Although a polyphyletic grouping of lophophorates was recovered in earlier molecular phylogenetic studies (Dunn *et al*., 2008; Hejnol *et al*., 2009; Hausdorf *et al*., 2010; Kocot *et al*., 2017), a monophyletic Lophophorata has gained support from more recent phylogenomic investigations (Nesnidal *et al*., 2013, 2014; Laumer *et al*., 2015, 2019; Marlétaz *et al*., 2019). Additionally, the interrelationships between the three phyla have been debated (Dunn *et al*., 2014; Carlson, 2016). Brachiopoda and Phoronida were suggested to share a last common ancestor by the Brachiozoa hypothesis (Dunn *et al*., 2008; Hejnol *et al*., 2009) or by including phoronids as members of brachiopods (Cohen and Weydmann, 2005; Carlson, 2007). The more recent molecular phylogenies, nevertheless, revealed a sister grouping of phoronids and bryozoans with brachiopods as the earliest branching group among lophophorates (Nesnidal *et al*., 2013, 2014; Laumer *et al*., 2015, 2019; Marlétaz *et al*., 2019). Despite these debates, brachiopods and phoronids are arguably closely related as they form a monophyletic or paraphyletic lineage under the alternative hypotheses of lophophorate phylogeny. Such a close evolutionary relationship between brachiopods and phoronids has been supported by an increasing number of morphological characters (e.g., Temereva and Tsitrin, 2015; Temereva and Kosevich, 2016; Temereva, 2017a, b).

The fossil record has provided the direct source of evidence for deciphering the origins of lophophorate body plans. The oldest putative total-group lophophorates can be traced back to the late Ediacaran skeletal *Namacalathus* (Zhuravlev *et al*., 2015), but the taxon was recently reinterpreted as a total-group lophotrochozoan (Shore *et al*., 2021). Among the three lophophorate phyla, typical brachiopods and bryozoans have biomineralized skeletons that are much more likely to be fossilized than the soft bodies of phoronids (Taylor *et al*., 2010). Brachiopods first appeared in the earliest Cambrian (Terreneuvian) and later became a major component of Paleozoic marine faunas (Carlson, 2007, 2016; Harper *et al*., 2017). Notably, the origin and early evolution of brachiopods have greatly advanced in the past two decades with contributions from the early Cambrian small shelly tommotiids (Holmer *et al*., 2008; Skovsted *et al*., 2008, 2009; Balthasar *et al*., 2009) and the soft-bodied preserved brachiopods of Cambrian Burgess Shale-type fossil deposits (Zhang *et al*., 2008, 2014; Balthasar and Butterfield, 2009). Bryozoans have long been thought to have their first appearance in the Early Ordovician Tremadocian (Taylor and Waeschenbach, 2015), but a very recent discovery traces their oldest fossil record back to the Cambrian Age 3 (Zhang *et al*., 2021). The fossil record of the soft-bodied phoronids is scattered, with putative Cambrian phoronids being controvertible (Chen and Zhou, 1997; Cohen and Weydmann, 2005; Conway Morris, 2006). Recently, the soft parts discovered in hyoliths suggest their lophophorate affinity (Moysiuk *et al*., 2017; Sun *et al*., 2018), but this remains a controversial issue (Liu *et al*., 2020; Smith, 2020; Li *et al*., 2020).

Here we probe deeper into the early fossil record of lophophorates by examining 68 new fossil specimens of *Conicula striata* Luo and Hu in Luo *et al*. (1999) preserved with exquisite soft parts from the early Cambrian Chengjiang Lagerstätte. *C. striata* exhibits a vermiform trunk with a pedicle and a U-shaped gut, as well as a lophophore surrounded by an undivided mantle. The mosaic morphologies of *C. striata* provide new insights into the origins of lophophorate phyla.

## Results

### Morphological description

*Conicula striata* possesses a conical and non-biomineralized body ranging from 13–71 mm in length (*n* = 29, mean = 28.6 mm, standard deviation = 11.9 mm; Figs. 1, 2i, 3a–e, 4a, 5 and Supplementary Figs. 1, 2; Supplementary Tables 1 and 2). An anatomical diagram of *C*. *striata* labelled with all morphological structures described is shown in Fig. 6a, and the interpretative drawings of most figured specimens are illustrated in Fig. 5. The body is divided into an upper lophophoral region and a lower trunk (‘lc’, ‘tr’ in Figs. 1, 2i, 3a–e, 4a, 5a–c, g, j–t and Supplementary Figs. 1, 2). The flaplike epistome present in extant lophophorates is unknown.

**Fig. 1.**
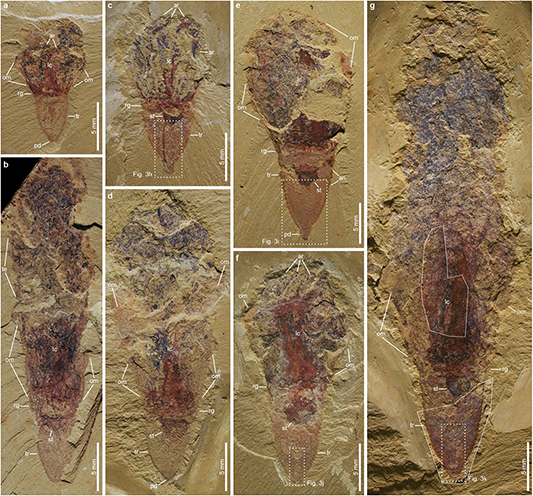
General morphology of *Conicula striata* in different sizes, with retracted or extended lophophores. (**a**, **c**, **e**, **f**) Specimens with retracted lophophores. **a** NIGP 171936A. **c** NIGP 171965. **e** NIGP 171938. **f** NIGP 171978. (**b**, **d**, **g**) Specimens with extended lophophores. **b** NIGP 171924. **d** NIGP 171977. **g** NIGP 171949. Note that the trunk has close size in (**a**, **b**), (**c**, **d**) and (**e**–**g**), respectively. Dotted boxes indicate areas of close-up images. Polygons indicate corresponding areas in the counterparts. an, annulation of trunk; ar, lophophoral arm; lc, lophophoral chamber; om, mantle; pd, pedicle; rg, medial ring between lophophoral chamber and trunk; st, stomach; te, tentacles; tr, trunk.

**Fig. 2.**
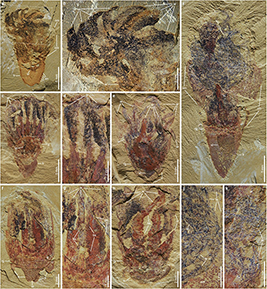
Lophophoral anatomy of *Conicula striata.* (**a**, **b**) NIGP 171920A. **a** Complete specimen. **b** Close-up view of coiled lophophore with lophophoral arms in black or brown. (**c**, **d**) NIGP 171948. **c** Complete specimen. **d** Close-up view of lophophoral arms with proximal part in red and distal part in black. **e** NIGP 171944A, showing at least 12 lophophoral arms in red or black. (**f**, **g**) NIGP 171957A. **f** Complete specimen. **g** Close-up view of lophophoral arms lined with feather-like bases of tentacles. **h** NIGP 171932B, showing lophophoral arms with bases of tentacles. (**i**, **j**) NIGP 171960. **i** Complete specimen. **j** Close-up view of tentacles. **k** NIGP 171939A, close-up view of lophophoral arms and tentacles in Fig. 3e. Dotted boxes indicate areas of close-up images. an, annulation of trunk; ar, lophophoral arm; bs, bases of tentacles; lc, lophophoral chamber; om, mantle; pd, pedicle; rg, medial ring between lophophoral chamber and trunk; st, stomach; te, tentacles; tr, trunk.

**Fig. 3.**
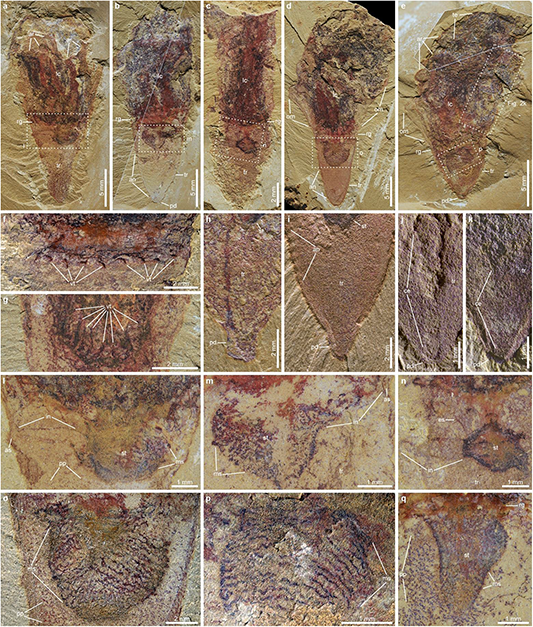
Trunk anatomy of *Conicula striata.* (**a**–**e**) Complete specimens. **a** NIGP 171956. **b** NIGP 171931. **c** NIGP 171981. **d** NIGP 171963. **e** NIGP 171939. (**f**, **g**) Close-up views of medial rings showing vascular tissues. **f** NIGP 171958A, area outlined in Supplementary Fig. 7e. **g** NIGP 176294A, area outlined in Supplementary Fig. 6e. (**h**, **i**) Close-up views of trunk with pedicle. **h** NIGP 171965, area outlined in Fig. 1c. **i** NIGP 171938, area outlined in Fig. 1e. (**j**, **k**) Close-up views of coelomic extension into lowermost trunk. **j** NIGP 171978, area outlined in Fig. 1f. **k** NIGP 171949, area outlined in Fig. 1g. (**l**, **m**) Close-up views of stomach with intestine reaching body wall of trunk. **l** NIGP 171956, area outlined in **a**. **m** NIGP 171931, area outlined in **b**. **n** NIGP 171981, close-up view of area outlined in **c** showing oesophagus, stomach and intestine. (**o**, **p**) Close-up views of musculature of stomach. **o** NIGP 171963, area outlined in **d**. **p** NIGP 171939, area outlined in **e**. **q** NIGP 171959, close-up view of area outlined in Fig. 4a showing stomach preserved in a different view from those in (**o**, **p**). Dotted boxes indicate areas of close-up images. Polygons indicate corresponding areas in the counterparts. an, annulation of trunk; ar, lophophoral arm; as, anus; bs, bases of tentacles; ce, coelomic lumen; es, oesophagus; in, intestine; lc, lophophoral chamber; ms, muscle of stomach; om, mantle; pd, pedicle; pp, putative papillae on trunk surface; rg, medial ring between lophophoral chamber and trunk; st, stomach; te, tentacles; tr, trunk; vt, vascular tissue.

**Fig. 4.**
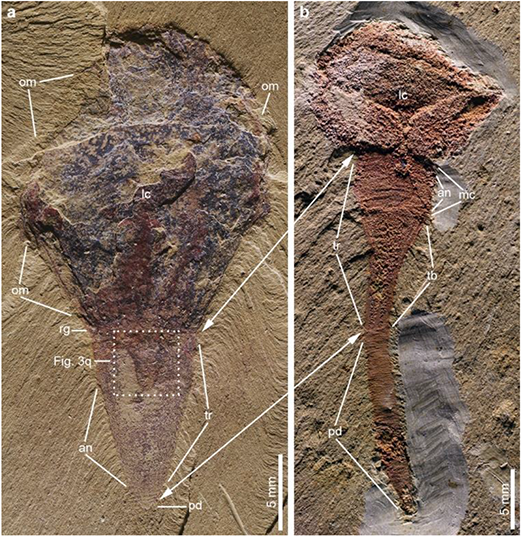
Comparative anatomy of *Conicula striata* and *Yuganotheca elegans*. **a** *Conicula striata*, NIGP 171959. **b** *Yuganotheca elegans*, NIGP 171987. Double-arrowed lines denote assumed homologous morphological boundaries of the two taxa. Dotted box indicates position of close-up image. an, annulation of trunk; lc, lophophoral chamber; mc, median collar; om, mantle; pd, pedicle; rg, medial ring between lophophoral chamber and trunk; tb, conical tube; tr, trunk.

**Fig. 5.**
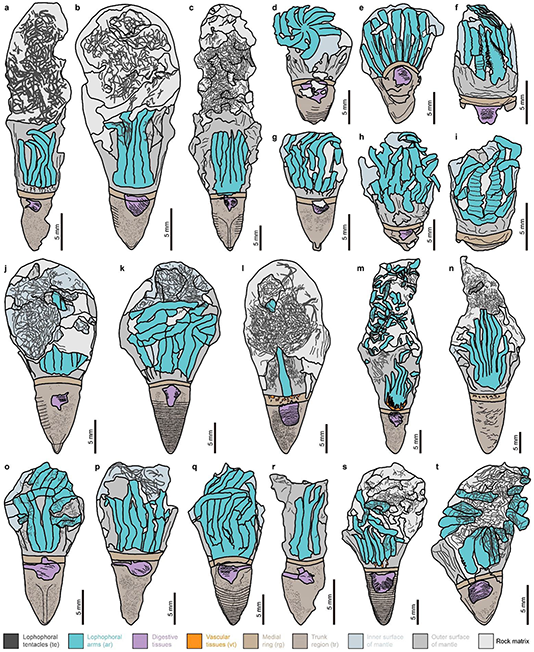
Interpretative drawings of selected specimens of *Conicula striata.* **a** NIGP 171924, as in Fig. 1b. **b** NIGP 171977, as in Fig. 1d. **c** NIGP 171949, as in Fig. 1g. **d** NIGP 171920A, as in Fig. 2a and Supplementary Fig. 4a. **e** NIGP 171948, as in Fig. 2c and Supplementary 5a. **f** NIGP 171957A, as in Fig. 2f and Supplementary Fig. 6a. **g** NIGP 171965, as in Fig. 1c. **h** NIGP 171944A, as in Fig. 2e and Supplementary Fig. 4i. **i** NIGP 171932B, as in Fig. 2h. **j** NIGP 171938, as in Fig. 1e. **k** NIGP 171959, as in Fig. 4a. **l** NIGP 171960, as in Fig. 1i. **m** NIGP 176294A, as in Fig. 3g and Supplementary Figs. 1b, 5e, part and counterpart combined. **n** NIGP 171958, as in Fig. 3f and Supplementary Fig. 6e, part and counterpart combined. **o** NIGP 171978, as in Fig. 1f. **p** NIGP 171956, as in Fig. 3a. **q** NIGP 171931, as in Fig. 3b. **r** NIGP 171981, as in Fig. 3c. **s** NIGP 171963, as in Fig. 3d and Supplementary Fig. 7e. **t** NIGP 171939, as in Fig. 3e. See legend for colours of different morphological structures.

**Fig. 6.**
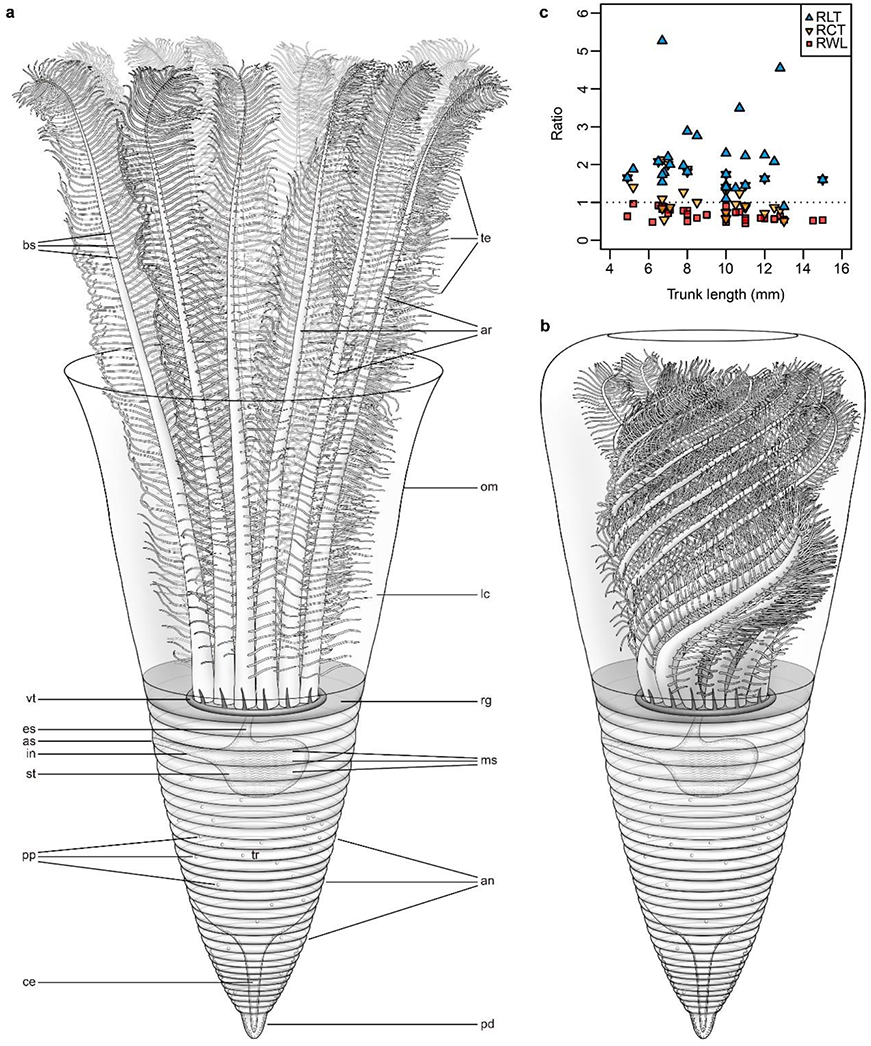
Reconstructions of *Conicula striata*. **a** Anatomical diagram with lophophore extended, showing labels of structures. **b** Anatomical diagram with lophophore retracted. **c** Scatterplot showing ratio of trunk width and length (RWL, red square), ratio of lophophoral chamber length and trunk length (RCT, orange downward triangle), and ratio of lophophoral region length and trunk length (RLT, blue upward triangle) relative to trunk length. See data in Supplementary Table 1. Mingjuan Feng made the drawings. Used with permission. an, annulation of trunk; ar, lophophoral arm; as, anus; bs, bases of tentacles; ce, coelomic lumen; es, oesophagus; in, intestine; lc, lophophoral chamber; ms, muscle of stomach; om, mantle; pd, pedicle; pp, putative papillae on trunk surface; rg, medial ring between lophophoral chamber and trunk; st, stomach; te, tentacles; tr, trunk; vt, vascular tissue.

The lophophoral region is 8–39 mm long and has an undivided saccular to tubular mantle that encases the lophophore (‘lc’ in Figs. 1, 2a, i, 3a–e, 4a, 5 and Supplementary Figs. 1–3; Supplementary Table 1). The mantle is flexible and is laterally expanded when the lophophore is retracted (‘om’ in Figs. 1a, c, e, 2a, i, 3d, e, 4a, 5d, g, j, l, s, t and Supplementary Figs. 1, 2a–c, 3). The lophophore is composed of at least 12 evenly spaced lophophoral arms in a circular arrangement (‘ar’ in Figs. 1c, f, 2b–h, 3a, 5), and it can extend up to 5X trunk length and to a maximum of 48 mm (‘lc’ in Figs. 1b, d, g, 5a–c, 6c and Supplementary Fig. 1). The lophophoral arms are erect or slightly curved in lateral view (‘ar’ in Figs. 1c, 2c, e, f, h, 3a, 5e–h), but one specimen preserved in dorsolateral view exhibits a counterclockwise spiral arrangement of the lophophoral arms (‘ar’ in Figs. 2a, b, 5d). These lophophoral arms are preserved in either reddish or dark colours under visible light (‘ar’ in Figs. 2b–h and 3a), but they have similar elemental profiles of iron, carbon, and phosphorus (Supplementary Figs. 4–7). Double rows of numerous evenly spaced tentacles are attached to each lophophoral arm in a pinnate arrangement (‘bs’, ‘te’ in Figs. 2f–k, 5f, i, l, t). The tentacles appear entwined with each other in lateral view (‘te’ in Figs. 1b, 2i–k, 3e, 5a–c, j–p, r–s). While the lophophoral arms have an average diameter of ∼1.0–1.5 mm (‘ar’ in Figs. 2b, d, g, h, 5), the tentacles measure ∼100–150 μm wide (‘te’ in Figs. 2g, j, k, 5a–c, j–p, r–s). No undisputed cilia could be identified on the tentacles because the preservation of such fine structures is rare.

Situated between the trunk and the lophophoral region is a narrow ring-shaped connective structure with clear upper and lower boundaries (‘rg’ in Figs. 1, 2c, f, i, 3a–e, 4a, 5 and Supplementary Figs. 1, 2a– c, 8). At least eight fine filamentous structures in a circular arrangement are associated with the medial ring, with each filament inserted to one lophophoral arm (‘vt’ in Figs. 3f, g, 5l–n). Elemental analysis reveals a distinct carbon-rich profile of these filaments (‘vt’ in Supplementary Figs. 5e–h and 6e–h), and these structures are interpreted as putative vascular tissues associated with lophophoral arms.

The trunk has a conical to bell-shaped outline (‘tr’ in Figs. 1, 2i, 3a–e, 4a, 5 and Supplementary Figs. 2, 3) and measures 5–15 mm long and 3–9 mm wide (*n* = 36, mean = 9.5 mm, standard deviation = 2.6 mm; Fig. 6c; Supplementary Table 1). Some specimens have as many as 25 annulations on the trunk, indicating that the body wall of the trunk is contractile (‘an’ in Figs. 1e, 2a, 3b, d, i, 4a, 5b–d, j, k, q, s and Supplementary Fig. 1a). A short pedicle is developed at the end of the trunk (‘pd’ in Figs. 1a, c–g, 3e, h– k, 4a, 5b, c, g, j, k, o, t and Supplementary Figs. 1c, 2), which is inserted by a narrow coelomic lumen emerging from the lower trunk (‘ce’ in Figs. 3j, k, 5c, o). On the trunk surface are papilla-like structures preserved as tiny reddish dots with diameters of < 100 µm (Figs. 3l, o, q, 5a–c, g, j, m, o–t).

The digestive tract is U-shaped and is differentiated into oesophagus, stomach, and intestine (Figs. 3l– n, 5o–r). The mouth opening, probably situated at the base of the lophophore, is not clearly exhibited on the specimens. The oesophagus is short and is connected dorsally to the stomach (‘es’ in Figs. 3n, 5r). The stomach is located within the upper part of the trunk, with its diameter ranging between 1–4 mm (‘st’ in Figs. 1b–g, 2a, c, f, i, 3a–e, 4a, 5a#x2013;h, j–m, o–t and Supplementary Figs. 1, 2). The outline of the stomach is generally globular or sac-like but varies with different angles of preservation (‘st’ in Fig. 3l– q). Equally spaced wavy musculature of the stomach is preserved (‘ms’ in Fig. 3l, m, o–q, 5a–c, e–h, j–m, o–q, s, t), showing positive elemental profiles of iron, carbon, and phosphorus (‘st’ in Supplementary Figs. 5a–d and 7). A roughly straight intestine emerges from the stomach almost horizontally or with a small angle and reaches the anus opened on the upper trunk (‘in’, ‘as’ in Figs. 3l–n, 5o–r).

**Fig. 7.**
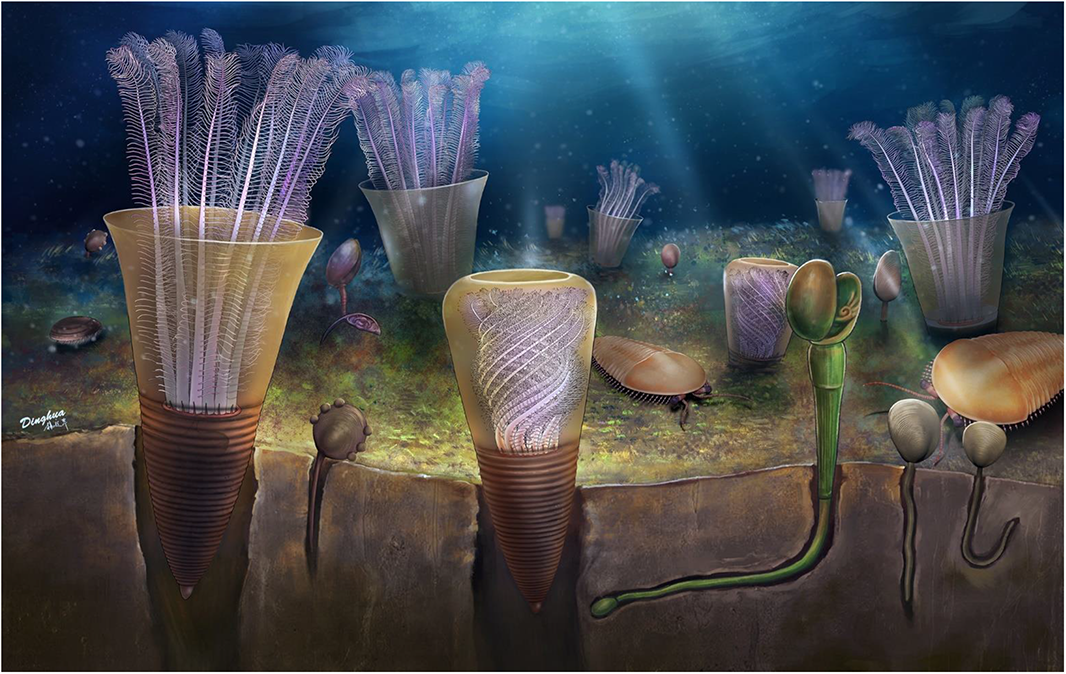
Ecological reconstruction of *Conicula striata*. Dinghua Yang made the drawing. Used with permission.

## Discussion

### Morphological comparisons

*Conicula striata* exhibits key morphological characteristics of lophophorates, including a lophophore, a sessile trunk and a U-shaped digestive tract (Fig. 6a). The lophophore of *C. striata* is functionally similar to that of bryozoans in being retractable (Hyman, 1959; Winston, 1977; Schwaha *et al*., 2020). However, bryozoans retract their lophophore into the body cavity, while *C. striata* keeps its retracted lophophore in a separate filtration chamber. Unlike bryozoans (Winston, 1977; Schwaha, 2020; Schwaha *et al*., 2020) and phoronids (Emig, 1982; Temereva, 2019), *C. striata* has its lophophore surrounded by a soft-bodied mantle as an extension of the body wall, which is the same state of the lophophore enclosed by the mantle lobes in brachiopods (Emig, 1992; Williams *et al*., 1997; Kuzmina *et al*., 2021). The at least 12 coiled lophophoral arms of *C. striata* are unique among lophophorates, although the double-row arrangement of tentacles on each lophophoral arm is shared by phoronids (Hyman, 1959; Emig, 1982; Brusca *et al*., 2016).

The vermiform trunk of *C. striata* is reminiscent of the phoronid body, but its conical morphology resembles the conical trunk of the stem-brachiopod *Yuganotheca elegans* (Zhang *et al*., 2014) (‘tr’, ‘tb’ in Fig. 4) rather than the generally cylindrical trunks of phoronids and bryozoans (Hyman, 1959; Winston, 1977; Emig, 1982; Schwaha, 2020). The trunk annulations of *C. striata* are absent in phoronids and bryozoans and are comparable to the annulations in the pedicles of *Y*. *elegans* and linguliform brachiopods (Zhang *et al*., 2008, 2014). Situated between the lophophoral region and the conical trunk, the medial ring of *C*. *striata* may be homologous to the median collar of *Y*. *elegans* (Zhang *et al*., 2014) (‘rg’, ‘mc’ in Fig. 4), although the former is more compacted and disc-shaped. The short pedicle of *C. striata* is associated with a central coelomic lumen, an extension of the coelomic cavity shared by the pedicles in *Y*. *elegans* and linguliform brachiopods (Zhang *et al*., 2008, 2014), which supports a homology between the pedicles in *C. striata* and stem-group brachiopods.

Among all metazoan phyla, a U-shaped gut is restricted to lophotrochozoans, including lophophorates, entoprocts, cycliophorans, sipunculans, and some molluscs, with the exception from pterobranchiate hemichordates (Hejnol and Martín-Durán, 2015; Budd and Jackson, 2016). Unlike the entoproctan condition with the anal opening falling inside the circle of tentacles (Zhang *et al*., 2013; Borisanova, 2018), the anus of *C*. *striata* is situated outside the tentacular apparatus and on the uppermost trunk part, an ectoproctan position akin to those in bryozoans and phoronids (Emig, 1982; Schwaha *et al*., 2020). Comparisons between the anuses of *C*. *striata* and brachiopods are not straightforward due to the variable gut morphologies among brachiopod lineages, such as the blind intestine in rhynchonelliforms, but a well-differentiated saccular stomach as in *C*. *striata* is shared by a number of extant brachiopod representatives including linguliforms, craniiforms, and rhynchonelliforms (Rudwick, 1970; McCammon, 1981; Nielsen, 1991; Williams and Carlson, 2007), which is not typical for bryozoans or phoronids (Emig, 1982; Schwaha *et al*., 2020).

The body plan of *C. striata* is distinct from the non-lophophorates bearing a U-shaped gut. Unlike the acoelomate cycliophorans without tentacles (Funch and Kristensen, 1995), *C. striata* is coelomate and possesses a typical tentacular apparatus. Although the tentacles of *C. striata* and pterobranchs share a feather-like organization, *C. striata* lack hemichordate characteristics such as a proboscis or mouth shield (Kaul-Strehlow and Röttinger, 2015). The absence of definite deuterostome features in *C. striata* shown here does not favour its affinity as a putative ambulacrarian closely related to the Cambrian problematics *Herpetogaster collinsi* and *Phlogites longus* (Caron *et al*., 2010). The general body plan of *C. striata* is much different in various aspects from sipunculans (Cutler, 1994) and molluscs (Wanninger and Wollesen, 2019), including tentacular organization and trunk shape.

### Functional morphology and ecology

As in all lophophorates, *Conicula striata* lacks effective locomotory organs such as appendages, hence it should be a sessile animal engaging in suspension feeding with its lophophore. In some specimens, the trunk is buried within the sediment, with the lophophoral region sticking out (Supplementary Fig. 1), supporting a semi-infaunal life style for *C*. *striata* (Fig. 7). This interpretation is consistent with the trunk morphologies such as annulations, a conical shape, and a pedicle, so the trunk would be capable in boring into shallow sediment. A sessile ecology is also compatible with a U-shaped gut morphology (Budd and Jackson, 2017). Because exposure of the anus would permit unhindered excretion from the intestine, for a feeding *C*. *striata* the maximum height of trunk buried in the sediments should be no higher than the anus on the upper trunk (Figs. 3a, b, l, m and 7). With a buried trunk and a retractable lophophore, *C*. *striata* could have protected its soft body from benthic and nektobenthic predators including various Cambrian arthropods (Erwin and Valentine, 2013), a survival strategy taken by extant phoronids and bryozoans (Winston, 1977; Emig, 1982). Although most specimens of *C*. *striata* are preserved individually, a couple of specimens show individuals living in aggregations (Supplementary Fig. 3d), like many other lophophorates (Hyman, 1959; Emig, 1982; Schwaha, 2020; Brusca *et al*., 2016).

The retraction and extension of lophophore of *C*. *striata* might be achieved by movements of associated retractor muscles and by hydraulic pressure from fluid in the coelomic or filtration cavity as in bryozoans (Winston, 1977; Schwaha, 2020; Schwaha *et al*., 2020). Although no convincing traces of these muscles have been found in the specimens, this is a common taphonomic phenomenon of rare preservation of muscle tissues in most Burgess Shale-type fossils (Gaines *et al*., 2008). The lophophoral architecture of *C*. *striata* comprising at least a dozen of lophophoral arms with numerous tentacles would greatly improve the efficiency of suspension feeding with more surface area for capturing minute food particles in the filtration cavity, as discussed in the relationships between feeding efficiency and lophophoral morphology (e.g., number of tentacles, size and shape of lophophore) in bryozoans (Winston, 1977). By comparing with other lophophorates, the double rows of tentacles may form a narrow food groove along the lophophoral arm where food particles are transported to the mouth (Hyman, 1959; Winston, 1977; Emig, 1982; Williams *et al*., 1997; Brusca *et al*., 2016). With its long extended lophophore, *C*. *striata* might also be capable of trapping relatively large food particles by folding their lophophoral tentacles over the prey and pulling it to the mouth, as indicated by the entwined tentacles in some specimens (Fig. 2j, k), analogous to some bryozoans (Winston, 1977; Schwaha *et al*., 2020). A potentially high efficiency of food ingestion was possibly accommodated by the wavy musculature of the stomach in *C*. *striata*, which indicates enhanced digestive abilities. In summary, the sophisticated and retractable lophophore of *C*. *striata* would make it an active suspension feeder and grant it effective defence under the intense ecological ‘arm race’ in the Cambrian origin of metazoan communities (Erwin and Valentine, 2013).

### Evolutionary implications

Various evolutionary scenarios on the origin of the brachiopod body plan have been proposed based on Cambrian fossils (Cohen *et al*., 2003; Carlson, 2007, 2016; Williams and Carlson, 2007; Zhang *et al*., 2014; Murdock *et al*., 2014; Harper *et al*., 2017). Among these, the brachiopod fold hypothesis of a *Halkieria*-like ancestor (Cohen *et al*., 2003) has not been favoured by a putative molluscan affinity of *Halkieria* (Vinther and Nielsen, 2005) and an articulated bivalved scleritome reconstruction of the tommotiid *Micrina* (Holmer *et al*., 2008, 2011). The tommotiids, problematic taxa with brachiopod-like scleritome features, have long been considered as a paraphyletic or polyphyletic assemblage linking phoronids and brachiopods (Holmer *et al*., 2008, 2011; Skovsted *et al*., 2008, 2009, 2011; Balthasar *et al*., 2009). This working hypothesis gained supports from recent phylogenetic reconstructions (Zhang *et al*., 2014; Sun *et al*., 2018). However, the precise phylogenetic affinities of tommotiids to total-group brachiopods remain debatable due to the paucity of morphological characters from skeletal fossils (Murdock *et al*., 2014), and the evolutionary insights from tommotiids have been blurred by the complicated history of biomineralization in lophophorates, with gains or losses of mineralized shells most likely occurring independently in different lineages (Taylor *et al*., 2010; Murdock, 2020). The soft anatomy of stem-group brachiopods, notably revealed by *Yuganotheca elegans*, has demonstrated the crucial roles of soft-bodied characters in deciphering the origin of brachiopods, which suggests a possibly unarmoured progenitor for brachiopods (Zhang *et al*., 2014). From such a soft-bodied perspective, the mosaic body of *C*. *striata* described above would provide novel and hopeful insights into the origin of the brachiopod body plan.

To resolve the phylogenetic position of *C*. *striata* among lophophorates, we conducted phylogenetic analyses based on a morphological data matrix of lophotrochozoans adapted from Sun *et al*. (2018). In parallel phylogenetic analyses with tommotiids either included or excluded, *C*. *striata* is resolved as an intermediate taxon between phoronids and brachiopods (Fig. 8 and Supplementary Fig. 9). Such a phylogenetic position of *C*. *striata* also remains consistent under various analytical methods and settings (Supplementary Table 4). The transitional phylogenetic placement of *C*. *striata*, with its chimeric body plan, uncovers several critical evolutionary transformations from a soft-bodied vermiform ancestor towards a bivalved body plan of brachiopods.

**Fig. 8.**
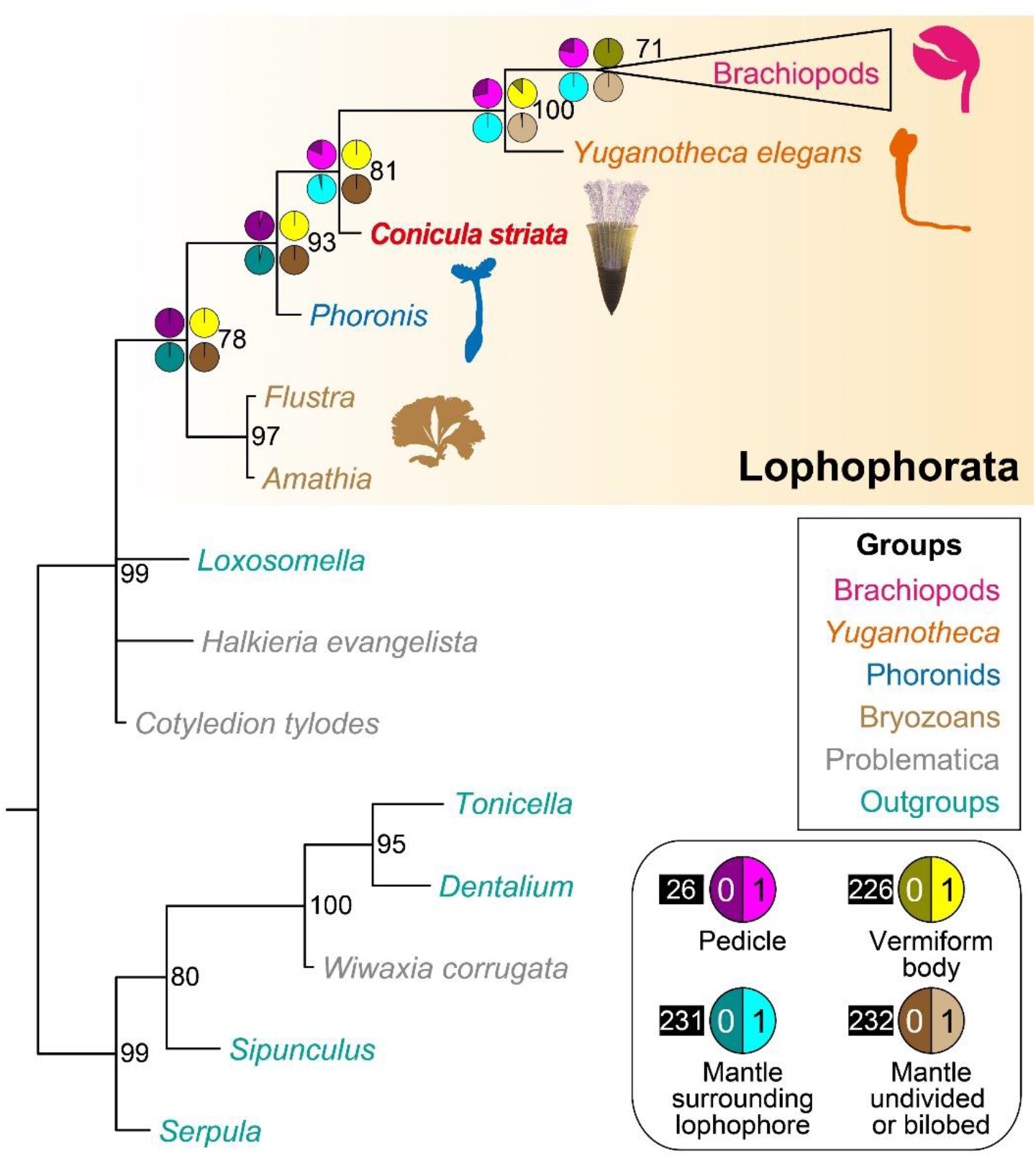
Phylogenetic placement of *Conicula striata* and early morphological evolution of Lophophorata. Lophophorate phylogeny resolved in a Bayesian analysis, showing *C*. *striata* in bold and red. The clade of brachiopods is collapsed. Nodal supports are posterior probabilities. See legends for colours of different taxonomic groups and characters. Pies indicate ancestral character states estimated from Bayesian inference in Supplementary Table 3. Indices of characters are indicated by numbers in black squares. See Supplementary Fig. 9a for a complete tree and Supplementary Fig. 9b for a tree with the problematic tommotiids and hyoliths included.

Phoronids and bryozoans do not possess a mantle covering the lophophore (Emig, 1982; Schwaha *et al*., 2020). *C*. *striata* bears an undivided mantle that surrounds the lophophore (Fig. 4a). In *Y*. *elegans*, there is an unmineralized bilobed mantle extension encasing the lophophore (Fig. 4b; Zhang *et al*., 2014). In brachiopods, a pair of biomineralized valves are secreted by the mantle lobes (Williams and Carlson, 2007). Ancestral character state reconstruction suggests that the mantle was an evolutionary novelty emerging in the deep origin of the brachiopod clade, and that the paired mantle lobes of brachiopods were derived from an undivided mantle in a phoronid-like progenitor as *C*. *striata* (Fig. 8; Supplementary Table 3). The unique coiled lophophore composed of at least 12 arms in *C*. *striata* also demonstrates the great morphological disparity of lophophores among Cambrian lophophorates (Zhang *et al*., 2003, 2008, 2009; Liu *et al*., 2020). The spiral arrangement of lophophoral arms in *C*. *striata* indicates that a single pair of lophophoral arms are a synapomorphy of total-group brachiopods including *Y*. *elegans*, and that the origination of paired lophophoral arms might be in concert with the appearance of bivalved body architecture during the origin of brachiopods (Zhang *et al*., 2014; Sun *et al*., 2018).

Phoronids possess a slender and tubular trunk (Emig, 1982), while *C. striata* and *Y. elegans* share a conical trunk part housing the visceral mass (Fig. 4; Zhang *et al*., 2014), and brachiopods lack a trunk that is well-separated from the lophophoral region (Williams *et al*., 1997). Such evolutionary transformations of trunk morphologies reinforce the hypothesis that a plesiomorphic vermiform trunk was successively reduced or fused with the lophophoral region in the deep evolutionary origin of brachiopods (Zhang *et al*., 2014), which is supported by the ancestral character state reconstruction (Fig. 8; Supplementary Table 3). Accompanying the reduction of the trunk, the brachiopod pedicle most likely originated from a primitive outgrowth of the body wall in their ancestors (Fig. 8; Supplementary Table 3), as illustrated by the short pedicle in *C*. *striata*, which exhibits a central coelomic lumen as in *Y*. *elegans* and linguliforms (Williams *et al*., 1997; Zhang *et al*., 2008, 2014).

In conclusion, *C*. *striata* is an extinct lophophorate taxon with a mosaic body plan, which reveals the crucial morphological transformations in a deep evolutionary root of brachiopods. It not only demonstrates the critical roles of soft-bodied fossils in resolving the early history of metazoans, but also highlights the tremendous morphological disparity of animals that emerged rapidly during the Cambrian explosion (Valentine, 2004; Erwin and Valentine, 2013).

## Methods

### Material studied

The 68 new specimens of *Conicula striata* examined in this study (NIGP 171920– 171986, 176294; Supplementary Tables 1 and 2), as well as a specimen of *Yuganotheca elegans* used for comparison (NIGP 171987), were collected from the main interval of the early Cambrian Chengjiang Lagerstätte within the Maotianshan Shale Member of the Yu’anshan Formation in the Haikou area, Kunming, Yunnan, China. Among these, NIGP 171920, 171921, 171985, and 171986 of *C*. *striata* were from the Mafang section, and all the other specimens came from the Jianshan section that is about 2 km away. Both the Mafang and Jianshan sections are close to the type locality of *C*. *striata* at the Ercaicun section (Luo *et al*., 1999), which is a few hundred metres away from Jianshan. A detailed geographic map of these fossil localities can be found in Zhao *et al*. (2012). All specimens are housed in the Nanjing Institute of Geology and Palaeontology, Chinese Academy of Sciences (NIGP). See Supplementary Table 2 for a full list of material examined.

### Image acquisition

The specimens were observed and photographed with a Zeiss Discovery V16 microscope or a Leica M125 microscope. A Nikon D810 camera fitted with a Nikon AF-S Micro NIKKOR 105 mm lens was used to photograph specimens in full view. Composite images integrating the part and counterpart of the same specimen were made by precise alignment in Adobe Photoshop CS6, where the overall tone, contrast, and brightness of photos were also adjusted. SEM-EDS elemental maps were obtained using an environmental scanning electron microscope (TESCAN MAIA3) equipped with an energy-dispersive X-ray spectrometer (Ultim Max, using AZtec software) under high vacuum conditions (< 0.001 Pa, 15 Kv) at the Nanjing Institute of Geology and Palaeontology, Chinese Academy of Sciences.

### Computed tomography

Tomographic scanning was carried out using a 3D X-ray microscope (Zeiss Xradia 520 Versa) at the micro-CT lab in the Nanjing Institute of Geology and Palaeontology (NIGP). For both the part and counterpart of NIGP 171957, a 0.4X object with LE2 filter was used to capture 1,800 projections over 180°, each with an exposure time of 2.5 s under voltage of 60 Kv and power of 5 W. Tomographic data were analyzed in VG StudioMax 3.0.

### Phylogenetic analysis

Our phylogenetic data matrix was adapted from the original dataset in Sun *et al*. (2018) by incorporating *Conicula striata* and related new characters with evolutionary implications (see character list in Supplementary Discussion). Two parallel analyses were run on the morphological matrix by either excluding or including tommotiids and hyoliths, resulting in 232 morphological characters with 40 or 55 taxa, respectively (Supplementary Data 1 and 2). Details on the data matrix analysed are described in the Supplementary Discussion. Tree reconstructions were conducted by using the Bayesian phylogenetic inference in MrBayes 3.2.7a (Ronquist *et al*., 2012). Two independent runs of 10,000,000 MCMC generations were performed, each containing four Markov chains under the Mkv + gamma model for discrete morphological character data (Lewis, 2001). In each run, trees were collected at a sampling frequency of every 1,000 generations, with the first 25% of sampled trees discarded as burn-in. The convergence of chains was confirmed by all effective sample sizes > 250 and was indicated by the ‘fuzzy caterpillar’-like traces in Tracer 1.7 (Rambaut *et al*., 2018). 50% majority-rule consensus trees of the remaining sampled trees were calculated and used as summaries of the results (Supplementary Fig. 9; Supplementary Data 1 and 2).

Analyses with different methods and various settings were conducted to test the robustness of the phylogenetic position of *C*. *striata*. Bayesian and parsimony methods were performed with MrBayes 3.2.7a (Ronquist *et al*., 2012) and TNT 1.5 (Goloboff and Catalano, 2016), respectively. Equal and implied weightings of characters were applied in parsimony analyses (Goloboff, 1993). The data matrix was treated with problematic taxa and new characters included or excluded. The results of these tests are summarized in Supplementary Table 4.

To investigate the origins of lophophorate body plans under the light of *C*. *striata*, ancestral character state reconstructions of characters 26, 226, 231, 232 were performed in MrBayes v.3.2.7a (Ronquist *et al*., 2012) with the same model parameters in phylogenetic tree reconstruction. Each character was treated as a separate partition from all the other characters. For each node of interest on the tree, probabilities of character states were calculated from posterior tree samples collected with a topological constraint at the node. The results are listed in Supplementary Table 3 and shown as pies in Fig. 8.

### Data availability

The authors declare that all data supporting the findings of this study are available within the paper and its supplementary information files. Specimens are deposited in the collections of the Nanjing Institute of Geology and Palaeontology, Chinese Academy of Sciences (prefix: NIGP). The source data underlying Fig. 8 and Supplementary Fig. 9 are provided in Supplementary Data 1 and 2.

## Acknowledgments

We thank Sandy Carlson and Doug Erwin for critical comments on earlier versions of the manuscript, Yan Fang for assistance in SEM-EDS analysis, Suping Wu for micro-CT scanning, and Mingjuan Feng and Dinghua Yang for artistic reconstruction. This study was supported by the National Key Research and Development Program of China (2021YFA0718100), the Strategic Priority Research Program (B) of the Chinese Academy of Sciences (XDB26000000), the National Natural Science Foundation of China (41872011, 41921002, 41902014, 42072006), and the State Key Laboratory of Palaeobiology and Stratigraphy, Nanjing Institute of Geology and Palaeontology, CAS (20192111).

## Author contributions

Han Zeng, Methodology, Formal analysis, Investigation, Visualization, Writing—original draft, Writing—review and editing; Xiangyuan Chen, Investigation, Visualization, Writing—original draft; Yao Liu, Investigation, Visualization, Writing—review and editing; Maoyan Zhu, Investigation, Writing— review and editing, Supervision, Funding Acquisition; Fangchen Zhao, Conceptualization, Investigation, Resources, Data Curation, Writing—review and editing, Supervision, Project Administration, Funding Acquisition; Aihua Yang, Conceptualization, Investigation, Resources, Data Curation, Writing—review and editing, Supervision, Project Administration, Funding Acquisition.

## Additional information

**Supplementary information** is available for this paper at https://doi.org/

**Correspondence** and requests for materials should be addressed to F.Z. or A.Y.

## Supplementary Information

**Supplementary Table 1.** Measurements of specimens of *Conicula striata*. Blue entries indicate the lophophores are retracted. Dark grey “NA” entries indicate valid measurements could not be made. Units are in millimetres. BL, body length; CL, lophophoral chamber length; LL, lophophoral region length; RCT, ratio of lophophoral chamber length and trunk length; RLT, ratio of lophophoral region length and trunk length; RWL, ratio of trunk width and length; TL, trunk length; TW, trunk width.

| Specimens | BL | TL | TW | CL | LL | RWL | RCT | RLT | Lophophoral status | Distortion |
| --- | --- | --- | --- | --- | --- | --- | --- | --- | --- | --- |
| NIGP 171920 | 17.0 | 6.7 | 5.5 | 7.3 | 10.3 | 0.82 | 1.09 | 1.54 | extended | normal |
| NIGP 171921 | NA | NA | 5.5 | 6.1 | 8.0 | NA | NA | NA | extended | normal |
| NIGP 171922 | NA | NA | 9.0 | 18.9 | 33.9 | NA | NA | NA | extended | twisted |
| NIGP 171923 | NA | 10.0 | 9.0 | NA | NA | 0.90 | NA | NA | extended | normal |
| NIGP 171924 | 32.0 | 8.5 | 5.0 | 8.5 | 23.5 | 0.59 | 1.00 | 2.76 | extended | normal |
| NIGP 171925 | 21.3 | 7.1 | 5.7 | 6.2 | 14.2 | 0.80 | 0.87 | 2.00 | extended | normal |
| NIGP 171926 | NA | NA | NA | NA | NA | NA | NA | NA | unknown | normal |
| NIGP 171927 | NA | NA | NA | NA | NA | NA | NA | NA | uncertain | twisted |
| NIGP 171928 | NA | NA | 9.5 | NA | NA | NA | NA | NA | extended | compacted |
| NIGP 171929 | NA | NA | NA | 15.9 | 15.9 | NA | NA | NA | retracted | normal |
| NIGP 171930 | NA | NA | 6.5 | 16.3 | 16.3 | NA | NA | NA | retracted | compacted |
| NIGP 171931 | 24.5 | 10.0 | 6.0 | 7.3 | 14.5 | 0.60 | 0.73 | 1.45 | uncertain | normal |
| NIGP 171932 | NA | NA | NA | NA | NA | NA | NA | NA | uncertain | twisted |
| NIGP 171933 | NA | NA | 6.2 | NA | NA | NA | NA | NA | unknown | compacted |
| NIGP 171934 | 20.0 | 7.0 | 6.0 | 13.0 | 13.0 | 0.86 | 1.86 | 1.86 | retracted | compacted |
| NIGP 171935 | NA | NA | 6.2 | NA | NA | NA | NA | NA | unknown | normal |
| NIGP 171936 | 13.0 | 4.9 | 3.1 | 8.1 | 8.1 | 0.63 | 1.65 | 1.65 | retracted | normal |
| NIGP 171937 | 21.0 | 10.0 | 4.8 | 5.7 | 11.0 | 0.48 | 0.57 | 1.10 | uncertain | normal |
| NIGP 171938 | 27.4 | 10.0 | 7.0 | 17.4 | 17.4 | 0.70 | 1.74 | 1.74 | retracted | normal |
| NIGP 171939 | 19.0 | 6.8 | 6.0 | 3.7 | 12.2 | 0.88 | 0.54 | 1.79 | extended | normal |
| NIGP 171940 | 19.5 | 6.5 | 6.0 | 13.5 | 13.5 | 0.92 | 2.08 | 2.08 | retracted | normal |
| NIGP 171941 | NA | 8.0 | 6.2 | NA | NA | 0.78 | NA | NA | extended | normal |
| NIGP 171942 | 48.0 | 10.7 | 8.0 | 13.3 | 37.3 | 0.75 | 1.24 | 3.49 | extended | normal |
| NIGP 171943 | NA | NA | 6.7 | 12.8 | 12.8 | NA | NA | NA | uncertain | normal |
| NIGP 171944 | NA | NA | 7.8 | NA | NA | NA | NA | NA | extended | compacted |
| NIGP 171945 | 21.5 | 7.8 | 6.1 | 9.9 | 15.4 | 0.78 | 1.27 | 1.97 | extended | normal |
| NIGP 171946 | NA | 6.5 | 6.0 | NA | NA | 0.92 | NA | NA | uncertain | compacted |
| NIGP 171947 | NA | 9.0 | 6.0 | NA | NA | 0.67 | NA | NA | uncertain | normal |
| NIGP 171948 | 18.3 | 6.7 | 6.0 | 5.6 | 11.6 | 0.90 | 0.84 | 1.73 | extended | normal |
| NIGP 171949 | 38.5 | 12.5 | 7.0 | 10.8 | 26.0 | 0.56 | 0.86 | 2.08 | extended | normal |
| NIGP 171950 | NA | NA | NA | NA | NA | NA | NA | NA | extended | normal |
| NIGP 171951 | 71.0 | 12.8 | 8.0 | 10.2 | 58.2 | 0.63 | 0.80 | 4.55 | extended | normal |
| NIGP 171952 | 42.0 | 6.7 | 6.0 | 14.3 | 35.3 | 0.90 | 2.13 | 5.27 | extended | normal |
| NIGP 171953 | NA | NA | 8.0 | NA | NA | NA | NA | NA | uncertain | normal |
| NIGP 171954 | NA | 14.5 | 7.5 | NA | NA | 0.52 | NA | NA | uncertain | normal |
| NIGP 171955 | 24.6 | 13.0 | 7.0 | 6.6 | 11.6 | 0.54 | 0.51 | 0.89 | extended | normal |
| NIGP 171956 | 25.0 | 10.5 | 7.8 | 10.0 | 14.5 | 0.74 | 0.95 | 1.38 | extended | normal |
| NIGP 171957 | NA | NA | 6.2 | NA | NA | NA | NA | NA | extended | normal |
| NIGP 171958 | NA | NA | 8.0 | NA | NA | NA | NA | NA | uncertain | normal |
| NIGP 171959 | 31.5 | 12.0 | 7.0 | 19.5 | 19.5 | 0.58 | 1.63 | 1.63 | retracted | normal |
| NIGP 171960 | 33.0 | 10.0 | 7.0 | 13.0 | 23.0 | 0.70 | 1.30 | 2.30 | extended | normal |
| NIGP 171961 | 24.0 | 10.0 | 6.0 | 14.0 | 14.0 | 0.60 | 1.40 | 1.40 | retracted | normal |
| NIGP 171962 | NA | NA | NA | NA | NA | NA | NA | NA | uncertain | compacted |
| NIGP 171963 | 39.0 | 12.0 | 7.0 | 8.5 | 27.0 | 0.58 | 0.71 | 2.25 | extended | normal |
| NIGP 171964 | NA | NA | NA | NA | NA | NA | NA | NA | uncertain | normal |
| NIGP 171965 | 15.0 | 5.2 | 5.0 | 7.3 | 9.8 | 0.96 | 1.40 | 1.88 | extended | normal |
| NIGP 171966 | NA | NA | NA | NA | NA | NA | NA | NA | extended | twisted |
| NIGP 171967 | NA | 6.2 | 3.0 | NA | NA | 0.48 | NA | NA | uncertain | normal |
| NIGP 171968 | NA | 11.8 | 7.0 | NA | NA | 0.59 | NA | NA | extended | normal |
| NIGP 171969 | NA | NA | NA | NA | NA | NA | NA | NA | retracted | compacted |
| NIGP 171970 | NA | NA | 7.1 | 20.0 | 20.0 | NA | NA | NA | retracted | normal |
| NIGP 171971 | NA | NA | NA | NA | NA | NA | NA | NA | uncertain | normal |
| NIGP 171972 | 21.0 | 7.0 | 5.0 | 14.0 | 15.4 | 0.71 | 2.00 | 2.20 | uncertain | normal |
| NIGP 171973 | 35.5 | 11.0 | 6.5 | 9.5 | 24.5 | 0.59 | 0.86 | 2.23 | extended | normal |
| NIGP 171974 | 39.0 | 15.0 | 8.0 | 24.0 | 24.0 | 0.53 | 1.60 | 1.60 | retracted | normal |
| NIGP 171975 | 22.5 | 8.0 | 4.0 | 14.5 | 14.5 | 0.50 | 1.81 | 1.81 | retracted | normal |
| NIGP 171976 | NA | NA | 6.2 | 20.0 | 40.0 | NA | NA | NA | extended | twisted |
| NIGP 171977 | 31.0 | 8.0 | 5.5 | 15.0 | 23.0 | 0.69 | 1.88 | 2.88 | extended | normal |
| NIGP 171978 | 27.0 | 11.0 | 5.0 | 16.0 | 16.0 | 0.45 | 1.45 | 1.45 | retracted | normal |
| NIGP 171979 | 27.0 | 11.0 | 5.8 | 10.0 | 16.0 | 0.53 | 0.91 | 1.45 | extended | normal |
| NIGP 171980 | NA | NA | NA | NA | NA | NA | NA | NA | uncertain | normal |
| NIGP 171981 | NA | NA | NA | NA | NA | NA | NA | NA | unknown | normal |
| NIGP 171982 | NA | NA | 5.7 | NA | NA | NA | NA | NA | uncertain | normal |
| NIGP 171983 | NA | 10.0 | 5.5 | NA | NA | 0.55 | NA | NA | uncertain | normal |
| NIGP 171984 | NA | NA | NA | NA | NA | NA | NA | NA | uncertain | normal |
| NIGP 171985 | NA | NA | NA | NA | NA | NA | NA | NA | uncertain | normal |
| NIGP 171986 | NA | NA | NA | NA | NA | NA | NA | NA | uncertain | normal |

**Supplementary Table 2.** Brief morphological description of specimens of *Conicula striata*.

| Specimens | Mantle | Tentacle | Trunk | Annulation | Pedicle | Stomach | Intestine | Medial ring | Vascular tissue | Completeness |
| --- | --- | --- | --- | --- | --- | --- | --- | --- | --- | --- |
| NIGP 171920 | + | + | + | + |  | + |  |  |  | Complete |
| NIGP 171921 | + | + |  |  |  |  |  |  |  | Incomplete |
| NIGP 171922 | + | + |  |  |  |  |  |  | + | Incomplete |
| NIGP 171923 | + | + | + |  | + | + |  | + |  | Nearly complete |
| NIGP 171924 | + | + | + |  | + | + |  | + |  | Complete |
| NIGP 171925 | + | + | + |  | + | + |  | + |  | Complete |
| NIGP 171926 |  |  | + | + | + |  |  |  |  | Incomplete |
| NIGP 171927 | + | + |  |  |  |  |  |  |  | Incomplete |
| NIGP 171928 | + | + | + |  | + | + |  | + |  | Complete |
| NIGP 171929 | + | + |  |  |  |  |  |  |  | Incomplete |
| NIGP 171930 | + | + | + |  |  | + |  | + |  | Complete |
| NIGP 171931 | + | + | + | + | + | + | + | + | + | Complete |
| NIGP 171932 |  | + |  |  |  |  |  |  |  | Incomplete |
| NIGP 171933 | + | + | + |  |  |  |  |  |  | Nearly complete |
| NIGP 171934 | + | + | + |  |  | + |  | + |  | Nearly complete |
| NIGP 171935 |  |  | + | + | + |  |  |  |  | Incomplete |
| NIGP 171936 | + | + | + |  | + |  |  | + |  | Near complete |
| NIGP 171937 | + | + | + | + | + |  |  | + |  | Complete |
| NIGP 171938 | + | + | + |  | + | + |  | + | + | Complete |
| NIGP 171939 | + | + | + |  | + | + |  | + |  | Complete |
| NIGP 171940 | + | + | + |  | + | + |  | + |  | Nearly complete |
| NIGP 171941 | + | + | + |  | + | + |  | + |  | Nearly complete |
| NIGP 171942 | + | + | + |  | + | + |  | + | + | Complete |
| NIGP 171943 | + | + |  |  |  |  |  | + |  | Incomplete |
| NIGP 171944 | + | + |  |  |  | + |  | + | + | Incomplete |
| NIGP 171945 | + | + | + |  | + | + |  | + |  | Complete |
| NIGP 171946 | + | + | + |  |  | + |  | + |  | Nearly complete |
| NIGP 171947 | + | + | + |  |  | + |  | + |  | Nearly complete |
| NIGP 171948 | + | + | + |  | + | + |  | + |  | Complete |
| NIGP 171949 | + | + | + | + |  | + |  | + |  | Complete |
| NIGP 171950 | + | + | + |  |  |  |  | + |  | Nearly complete |
| NIGP 171951 | + | + | + |  |  | + |  | + |  | Complete |
| NIGP 171952 | + | + | + |  | + | + |  | + | + | Complete |
| NIGP 171953 | + | + | + |  |  | + |  | + |  | Incomplete |
| NIGP 171954 | + | + | + |  |  |  |  | + |  | Incomplete |
| NIGP 171955 | + | + | + |  |  |  |  | + |  | Nearly complete |
| NIGP 171956 | + | + | + | + | + | + | + | + |  | Nearly complete |
| NIGP 171957 | + | + |  |  |  |  |  |  |  | Incomplete |
| NIGP 171958 | + | + | + |  |  |  |  | + | + | Incomplete |
| NIGP 171959 | + | + | + | + | + |  |  | + |  | Nearly complete |
| NIGP 171960 | + | + | + |  | + | + |  | + | + | Complete |
| NIGP 171961 | + | + | + | + | + | + |  | + |  | Complete |
| NIGP 171962 | + | + | + |  |  | + |  |  |  | Complete |
| NIGP 171963 | + | + | + | + |  | + |  | + | + | Complete |
| NIGP 171964 | + | + |  |  |  |  |  |  |  | Incomplete |
| NIGP 171965 | + | + | + |  | + | + |  | + |  | Nearly complete |
| NIGP 171966 |  | + |  |  |  |  |  |  |  | Incomplete |
| NIGP 171967 | + | + | + |  | + |  |  | + |  | Incomplete |
| NIGP 171968 | + | + | + |  | + |  |  | + |  | Nearly complete |
| NIGP 171969 | + | + | + |  |  | + |  | + |  | Complete |
| NIGP 171970 | + | + | + |  | + | + |  | + |  | Incomplete |
| NIGP 171971 | + | + |  |  |  |  |  |  |  | Incomplete |
| NIGP 171972 | + | + | + |  | + | + |  | + |  | Nearly complete |
| NIGP 171973 | + | + | + |  | + | + |  | + | + | Complete |
| NIGP 171974 | + | + | + |  | + | + |  | + | + | Complete |
| NIGP 171975 | + | + | + |  | + |  |  | + |  | Complete |
| NIGP 171976 | + | + |  |  |  |  |  | + |  | Incomplete |
| NIGP 171977 | + | + | + |  | + | + |  | + | + | Complete |
| NIGP 171978 | + | + | + |  |  | + |  | + |  | Nearly complete |
| NIGP 171979 | + | + | + |  | + | + |  | + |  | Nearly complete |
| NIGP 171980 | + | + | + |  |  |  |  | + |  | Nearly complete |
| NIGP 171981 | + | + | + |  |  | + | + | + |  | Incomplete |
| NIGP 171982 | + | + |  |  |  |  |  |  |  | Incomplete |
| NIGP 171983 | + |  | + |  | + | + |  | + |  | Incomplete |
| NIGP 171984 | + | + |  |  |  |  |  | + |  | Incomplete |
| NIGP 171985 | + |  |  |  |  |  |  |  |  | Incomplete |
| NIGP 171986 | + |  | + |  | + |  |  |  |  | Incomplete |
| NIGP 176294 | + | + | + |  |  | + |  | + | + | Nearly complete |

**Supplementary Table 3.**
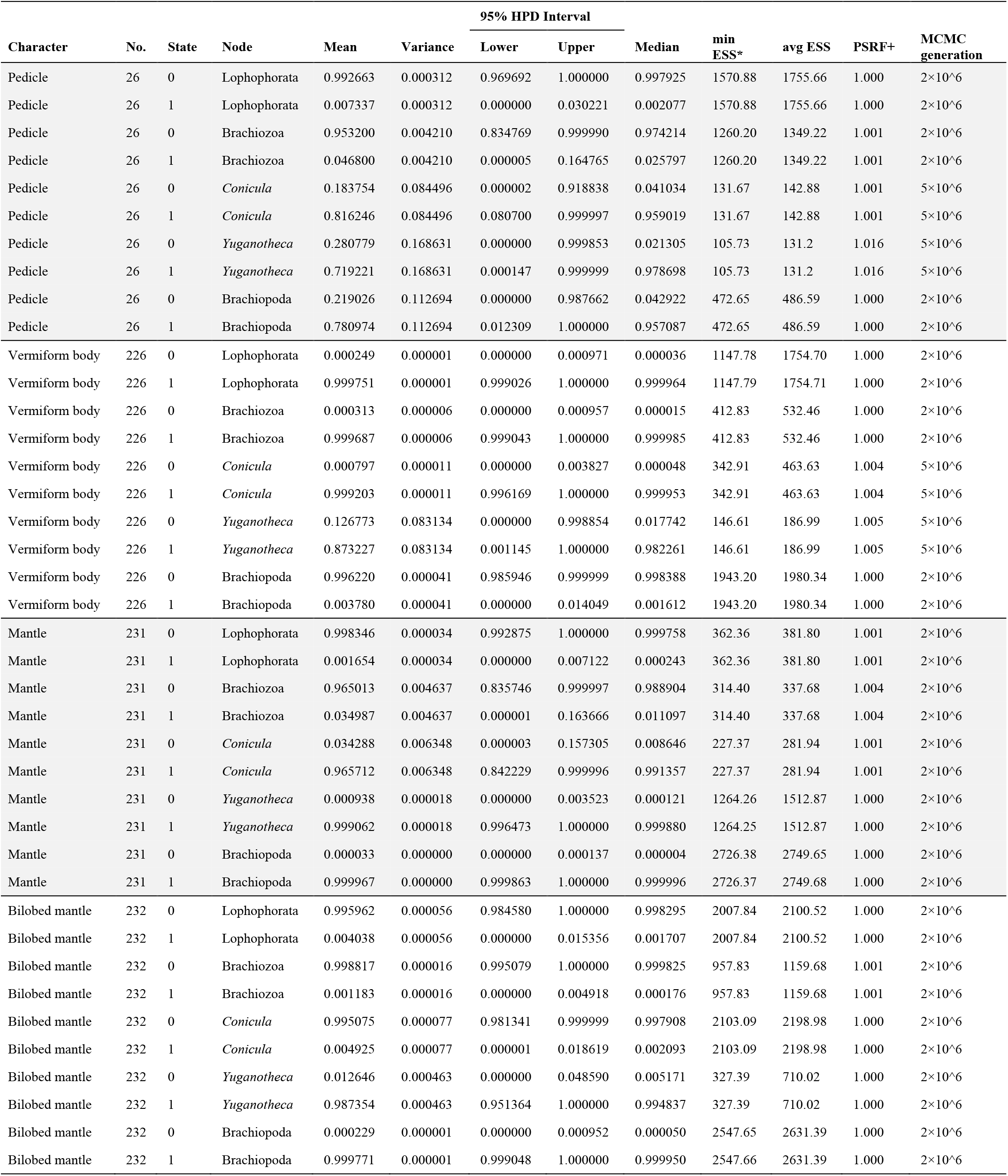
Bayesian inference of ancestral character states from MrBayes output. Mean probabilities of character states are illustrated as pies in Fig. 8. Node under estimation is indicated by the name of taxon. MCMC generation stands for number of Markov chain Monte Carlo (MCMC) generations with the first 25% samples as burn-in. ESS, Effective Sample Size; HPD, Highest Posterior Density; PSRF, Potential Scale Reduction Factor.

**Supplementary Table 4.** Phylogenetic position of *Conicula striata* under tests with different methods and settings. The methods applied were Bayesian and parsimony methods using MrBayes and TNT, respectively. The parsimony analyses involved equal and implied weighting of characters. The data matrix was treated with problematic taxa and new characters included or excluded. See notes for the topological position of problematic taxa.

| Method | Concavity constant *k* | Taxa | Characters | Position of *Conicula striata* | Notes |
| --- | --- | --- | --- | --- | --- |
| Bayesian | NA | All taxa | All characters | Between Phoronida and (Brachiopoda + Hyolitha) |  |
| Bayesian | NA | No problematic taxa | All characters | Between Phoronida and Brachiopoda |  |
| Bayesian | NA | All taxa | No new characters | In a big polytomy with Phoronida, (Brachiopoda + Hyolitha), Bryozoa, Entoprocta, and more. |  |
| Bayesian | NA | No problematic taxa | No new characters | In a polytomy with Phoronida and Brachiopoda. |  |
| Parsimony | 2, 3, 5, 10, ∞ | All taxa | All characters | Between Phoronida and (Brachiopoda + Hyolitha) | *Namacalathus* is topologically unstable with $k = \infty$. |
| Parsimony | 2, 3, 5, 10, ∞ | No problematic taxa | All characters | Between Phoronida and Brachiopoda |  |
| Parsimony | 2, 3, 5, 10, ∞ | All taxa | No new characters | Between Phoronida and ((Brachiopoda + Hyolitha) + *Namacalathus*) | The position of the problematic *Namacalathus* is questionable. |
| Parsimony | 2, 3, 5, 10, ∞ | No problematic taxa | No new characters | Between Phoronida and Brachiopoda | *Halkieria* is topologically unstable with $k = \infty$. |

**Supplementary Fig. 1.**
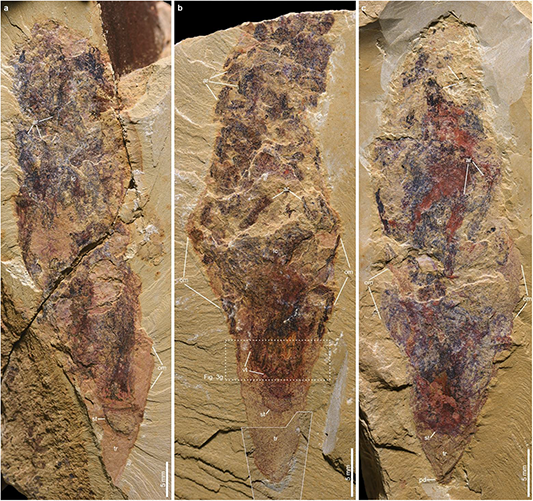
Additional specimens of *Conicula striata* showing extended lophophores. **a** NIGP 171951. **b** NIGP 176294. **c** NIGP 171952. Dotted box indicates areas of close-up images. Polygons indicate corresponding areas in the counterparts. ar, lophophoral arm; lc, lophophoral chamber; om, mantle; pd, pedicle; st, stomach; tr, trunk.

**Supplementary Fig. 2.**
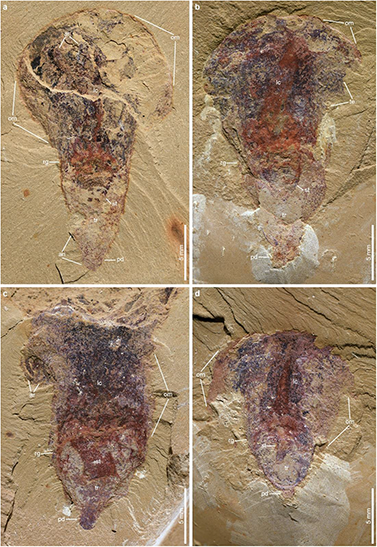
Additional specimens of *Conicula striata* showing pedicle. **a** NIGP 171961. **b** NIGP 171940. **c** NIGP 171923. **d** NIGP 171928. an, annulation of trunk; ar, lophophoral arm; lc, lophophoral chamber; om, mantle; pd, pedicle; rg, medial ring between lophophoral chamber and trunk; st, stomach; tr, trunk.

**Supplementary Fig. 3.**
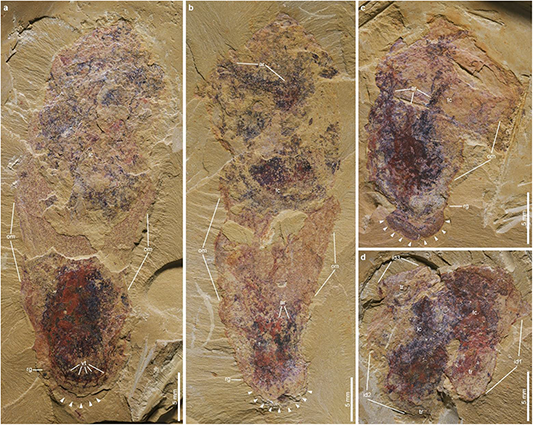
Specimens showing life habits of *Conicula striata*. (**a**–**c**) Specimens with trunk embedded within the sediments. **a** NIGP 171922. **b** NIGP 171976. **c** NIGP 171930. **d** Three closely preserved specimens, NIGP 171980A–C. Arrowheads denote the boundary between non-embedded and embedded body parts along the water-sediment surface. ar, lophophoral arm; id1–3, individuals of *C*. *striata* 1–3; lc, lophophoral chamber; om, mantle; rg, medial ring between lophophoral chamber and trunk; st, stomach; tr, trunk.

**Supplementary Fig. 4.**
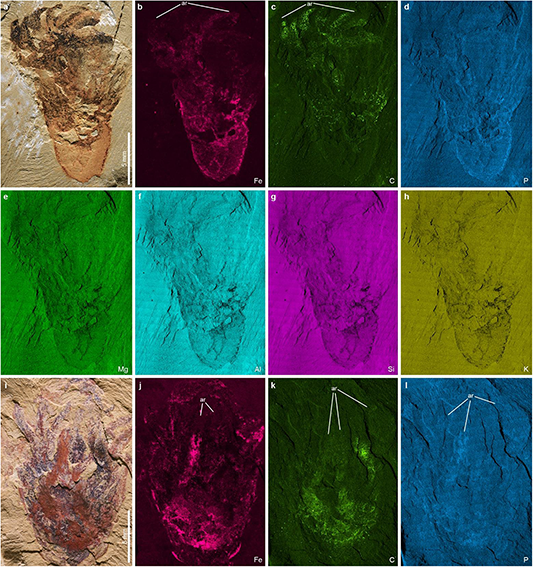
SEM-EDS elemental maps of *Conicula striata*. (**a**–**h**) NIGP 171920. (**i**–**l**) NIGP 171944. Note that the lophophoral tentacles preserved as positive iron and carbon imprints that are little overlapped with each other. ar, lophophoral arm.

**Supplementary Fig. 5.**
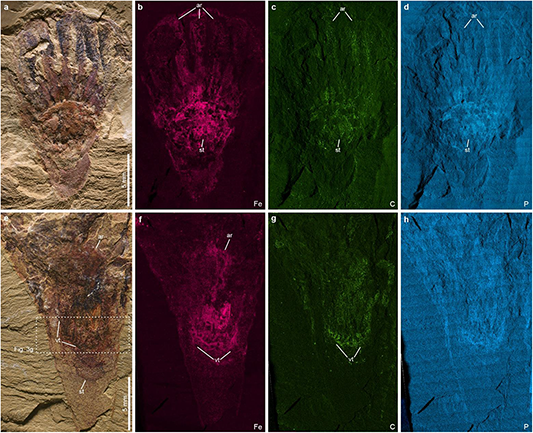
SEM-EDS elemental maps of *Conicula striata*. (**a**–**d**) NIGP 171948. Note that the lophophoral tentacles have similar positive profiles of iron, carbon, and phosphorus, despite the colour differences in proximal and distal portions under visible light. (**e**–**h**) NIGP 171949. Note that the putative vascular tissues are preserved as iron and carbon imprints. Dotted boxes indicate areas of close-up images. ar, lophophoral arm; st, stomach; vt, vascular tissue.

**Supplementary Fig. 6.**
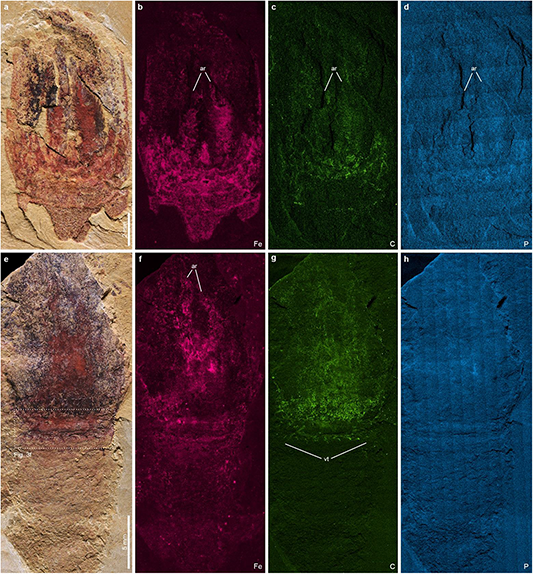
SEM-EDS elemental maps of *Conicula striata*. (**a**–**d**) NIGP 171957. Note that the lophophoral tentacles preserved as positive iron, carbon, and phosphorus imprints. (**e**–**h**) NIGP 171958A. Note that the putative vascular tissues are mainly preserved as carbon imprints. Dotted boxes indicate areas of close-up images. ar, lophophoral arm; vt, vascular tissue.

**Supplementary Fig. 7.**
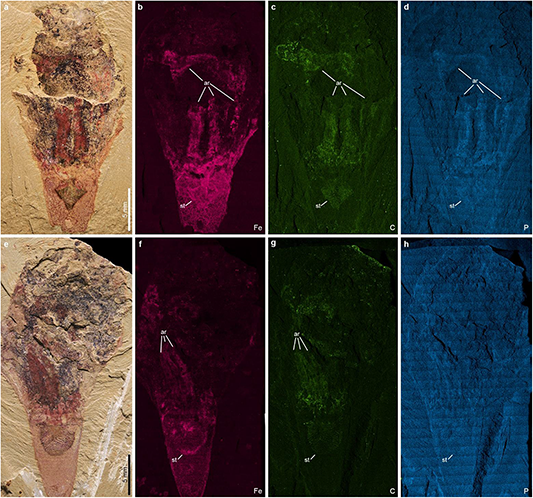
SEM-EDS elemental maps of *Conicula striata*. (**a**–**d**) NIGP 171970. (**e**–**h**) NIGP 171963. Note that the lophophoral tentacle and stomach are preserved with positive profiles of iron, carbon, and phosphorus. ar, lophophoral arm; st, stomach.

**Supplementary Fig. 8.**
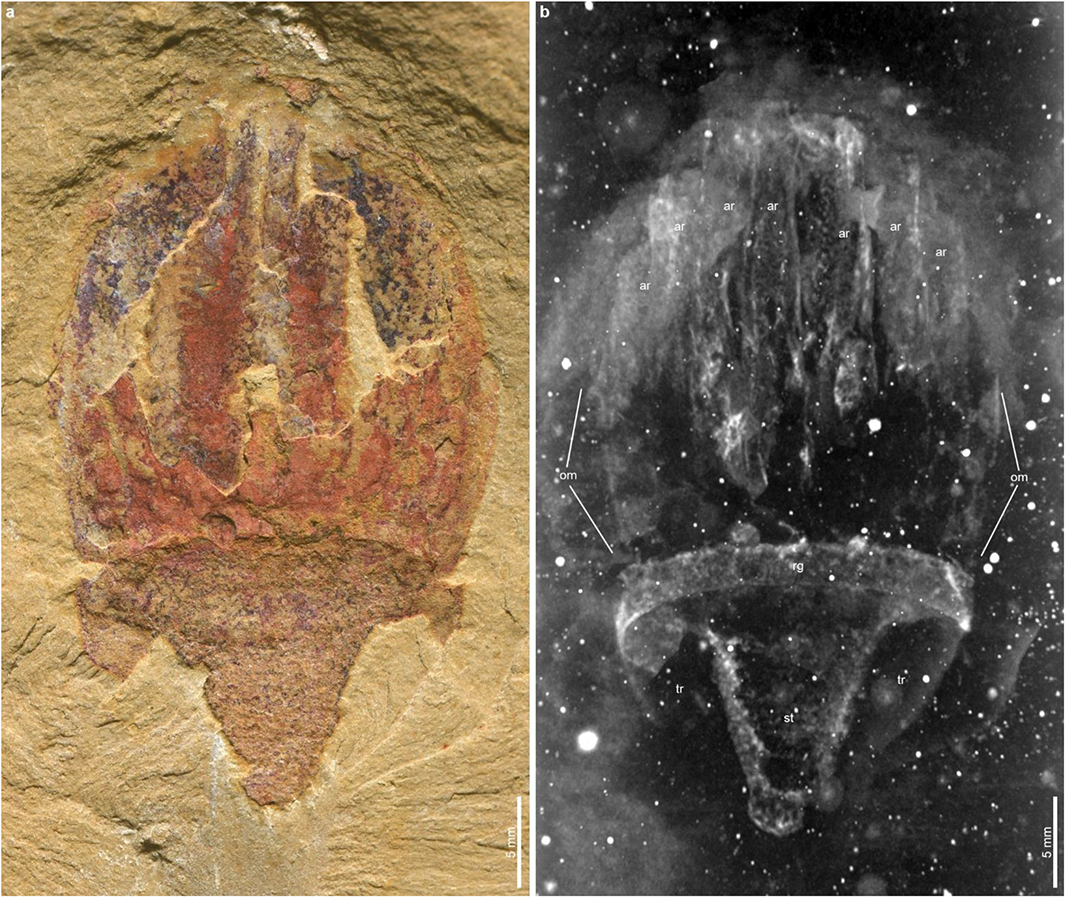
X-ray computed tomography of *Conicula striata*, NIGP 171957B. (**a**) Photo under normal light. (**b**) Tomographic image in maximum projection mode. ar, lophophoral arm; om, mantle; rg, medial ring between lophophoral chamber and trunk; st, stomach; tr, trunk.

**Supplementary Fig. 9.**
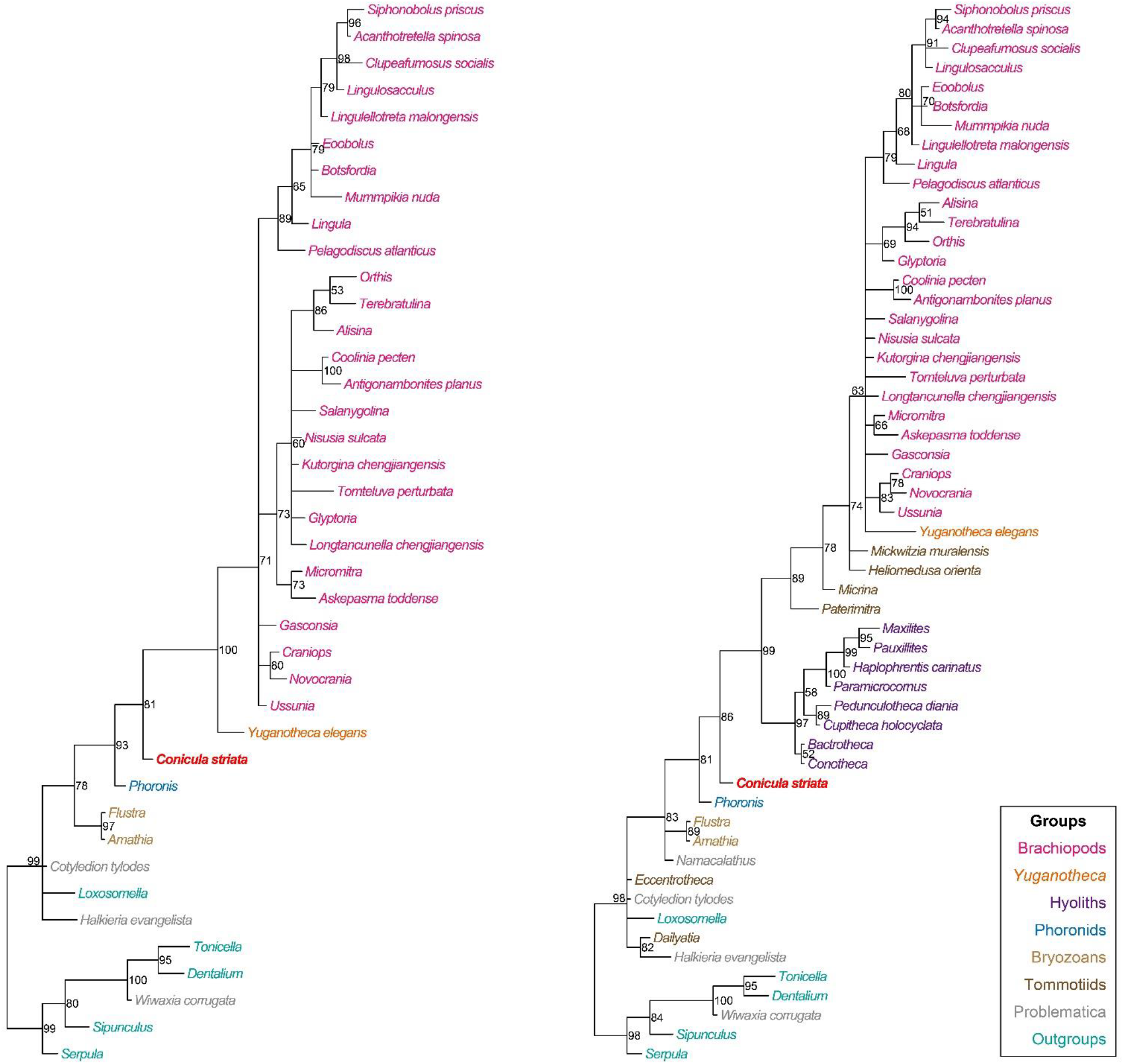
Detailed phylogenetic contexts of *Conicula striata*. **a** Tommotiids and hyoliths included. **b** Tommotiids and hyoliths excluded. Phylogenetic reconstructions are from Bayesian analyses. Nodal supports are posterior probabilities. See legends for colours of different taxonomic groups.

### Supplementary Discussion

#### Systematic palaeontology

Lophotrochozoa Halanych *et al*., 1995

Lophophorata Hyman, 1959

*Conicula striata* Luo and Hu in Luo *et al*. 1999

(Figs. 1–3, 4a and Supplementary Figs. 1–8)

*1999 *Conicula striata* Luo *et al*., p. 87, text-fig. 35, pl. 22, fig. 5a.

.2002 *Conicula striata* Chen *et al*., pl. 20, fig. 2.

**Holotype.** Part He-f-6-5-112 and counterpart He-f-6-5-113.

**Other material**. NIGP 171920–171986, among which the figured specimens in this paper include NIGP 171920, 171922, 171924, 171931, 171932, 171936, 171938, 171939, 171944, 171948, 171949, 171956–171960, 171963, 171965, 171970, 171977, 171978 and 171981.

**Emended diagnosis.** Lophophoral region enclosed by an undivided mantle. Lophophore retractable, composed of at least 12 lophophoral arms and numerous tentacles. Conical trunk possessing striated annulations and bearing a terminal pedicle. U-shaped gut differentiated, comprising an inflated stomach and an intestine opened at upper part of trunk.

**Description.** See the main text.

**Remarks.** Before the current study, *C*. *striata* was only known from the single holotype specimen and was identified as a lophophorate with a very brief description in the Chinese literature(Luo *et al*., 1999; Chen *et al*., 2002). The original anatomical description includes a conical body bearing horizontal striation, a trumpet-shaped cavity and eight slightly curved short tentacles (Luo *et al*., 1999). The holotype specimen was later illustrated again in colour but was not further described (Chen *et al*., 2002). The morphological description of *C*. *striata* is extensively updated in the main text of this paper.

**Occurrence.** Ercaicun, Jianshan, and Mafang sections, Haikou, Kunming, Yunnan. Maotianshan Shale Member, Yu’anshan Formation. *Wutingaspis*–*Eoredlichia* Interval Zone, Cambrian Series 2, Stage 3.

#### Phylogenetic analysis

*Conicula striata* and associated new characters of evolutionary significance were incorporated into the original morphological data matrix in Sun *et al*. (2018), which sampled representative fossil and extant lophophorates and other notable lophotrochozoans. To accommodate the anatomical information from *C*. *striata*, the coding of characters 28, 120 and 122 for *Yuganotheca elegans* in Sun *et al*. (2018) has been changed as 2→1, 1→? and 1→?, respectively. For character 28, the biomineralization of pedicle in *Y. elegans* should be absent if its conical tube is not considered as part of the pedicle. For characters 120 and 122, we consider the states of umbonal perforation in the ventral valve of *Y. elegans* are questionable because the connection between its specialized median collar and valve is unclear. To investigate the impacts of phylogenetically questionable tommotiids on tree topology (e.g., Mudrock *et al*., 2014), two parallel analyses with tommotiids and hyoliths excluded (41 taxa in total) or included (55 taxa in total) were performed (Supplementary Data 1 and 2). The phylogenetic position of *C*. *striata* is consistently retrieved as an intermediate taxon under both settings (Fig. 8 and Supplementary Fig. 9).

#### Character list

The characters 1–225 below were from Sun *et al*. (2018) (see data deposited at MorphoBank, project number 2800: http://morphobank.org/permalink/?P2800). Seven new morphological characters were added to the phylogenetic dataset of Sun *et al*. (2018) to test the potential homologies of structures revealed by *C*. *striata*, resulting into a total of 232 characters (Supplementary Data 1 and 2). These new characters are listed at the end of the character list below.

1. Brephic shell: Embryonic shell. Absent (0); Present (1).
2. Brephic shell: Morphology. [Transformational character] (0); Flat, disc-like (cf. Micrina) (1); Three prominent lobes forming a Y (cf. Paterimitra) (2); Spherical (3); Fusiform (4).
3. Brephic shell: Embryonic shell extended in larvae. [Transformational character] (0); Not extended; embryonic shell contiguous with adult shell (1); Extended into larval shell, separated from adult shell by prominent nick (2).
4. Brephic shell: Surface ornament. [Transformational character] (0); Smooth (1); Rounded pits (2); Polygonal impressions (3); Pustulose (4).
5. Brephic shell: Larval attachment structure. [Transformational character] (0); Without evidence of pedicle (1); With evidence of pedicle (2).
6. Brephic shell: Setulose No evidence of setae in embryonic shell. (0); Setae present (1).
7. Brephic shell: Setal sacs. Absent (0); Present (1).
8. Brephic shell: Setal sacs: Number. [Transformational character] (0); One pair (1); Two pairs (2); Three pairs (3).
9. Larval chaetae: Paired bundles. Absent (0); Present (1).
10. Adult setae: Presence. Absent (0); Present (1).
11. Adult setae: Secretion. [Transformational character] (0); By basal microvilli (1); Epicuticular (2).
12. Adult setae: Secretion: Microvillar diameter. [Transformational character] (0); Uniform (1); Decreasing towards seta margin (2).
13. Adult setae: Secretion: Microvillar canal aspect. [Transformational character] (0); Round (1); Polygonal; close-packed (2).
14. Adult setae: Composition. [Transformational character] (0); Chitin (1); Horny protein (2).
15. Adult setae: Enamel. Absent (0); Present (1).
16. Adult setae: Distribution. [Transformational character] (0); Uniform(1); Only present at margins of shell (2); In bundles, repeated on each metamere if serial repetition present (3); In digestive tract (4).
17. Adult setae: Constitution. [Transformational character] (0); Solid, blade-like (1); Basal invagination (2).
18. Adult setae: Projecting knobs. Absent (0); Individual peripheral projections present (1).
19. Adult setae: Circling coronae. Absent (0); Present (1).
20. Body organization: Serial repetition. Absent (0); Present (1).
21. Body organization: Foot. Absent (0); Present (1).
22. Body organization: Coelom. Absent: adults acoelomate (0); Present: true coelomic cavities differentiated (1).
23. Body organization: Coelomoducts: Number. [Transformational character] (0); Single (1); Multiple (2).
24. Body organization: Gills. Absent (0); Present (1).
25. Body organization: Circulatory system. [Transformational character] (0); Epithelially lined (1); Poorly defined, involving sinuses and lacunae (2); Closed circulatory system (3).
26. Pedicle: Presence. Absent (0); Present (1).
27. Pedicle: Constitution. [Transformational character] (0); Massive or uniform (1); Densely stacked tabular discs (2).
28. Pedicle: Biomineralization. [Transformational character] (0); Absent (1); Present (2).
29. Pedicle: Bulb. Absent (0); Present (1).
30. Pedicle: Distal rootlets. Absent (0); Present (1).
31. Pedicle: Tapering. [Transformational character] (0); Uniform thickness (1); Tapering (2).
32. Pedicle: Coelomic region. [Transformational character] (0); Absent (1); Present (2).
33. Pedicle: Surface ornament. [Transformational character] (0); Smooth (1); Regular annulations (2); Irregular wrinkles (3).
34. Pedicle: Nerve impression. Absent (0); Present (1).
35. Mantle canals: Presence. Absent (0); Present (1).
36. Mantle canals: Morphology. [Transformational character] (0); Pinnate (=lemniscate) (1); Bifurcate (2); Baculate (3); Saccate (4).
37. Mantle canals: vascula lateralia. Absent (0); Present (1).
38. Mantle canals: vascula media. Absent (0); Present (in dorsal valve) (1).
39. Mantle canals: vascula terminalia. [Transformational character] (0); Exclusively marginal (peripheral) (1); Directed peripherally and (intero)medially (2).
40. Perioral tentacular apparatus: Presence. Absent (0); Present (1).
41. Perioral tentacular apparatus: Origin. [Transformational character] (0); Prostomium (i.e. anterior of larval prototroch) (1); Second pair of coelomic sacs, at metamorphosis (2); Mid-trunk, prior to metamorphosis (3).
42. Perioral tentacular apparatus: Tentacle disposition. [Transformational character] (0); Single side (1); Both sides (2).
43. Perioral tentacular apparatus: Tentacle rows per side in trocholophe stage. No additional ablabial row (0); Adlabial and ablabial row (1).
44. Perioral tentacular apparatus: Tentacle rows per side in post-trocholophe stage. No additional ablabial row (0); Adlabial and ablabial row (1).
45. Perioral tentacular apparatus: Median tentacle in early development. Absent (0); Present (1).
46. Perioral tentacular apparatus: Site of tentacle addition. [Transformational character] (0); At two points located behind the mouth (1); At the tip of each lophophore arm (2).
47. Perioral tentacular apparatus: Innervation. [Transformational character] (0); Extensions of a circumoral nerve ring (1); Nerve rings within the tentacle ring itself (2).
48. Perioral tentacular apparatus: Inner nerve ring. Not reduced (whether present or absent due to absence of lophophore nerve rings) (0); Reduced, weakly developed or absent in adults (1).
49. Perioral tentacular apparatus: Outer nerve ring. Not reduced (whether present or absent due to absence of lophophore nerve rings) (0); Reduced, weakly developed or absent in adults (1).
50. Perioral tentacular apparatus: Musculature. [Transformational character] (0); Outer main tentacle muscle; two pairs of inner longitudinal muscles (1); Peripheral muscle fibres (2); Core of longitudinal muscle fibres surrounded by circular muscles (3).
51. Perioral tentacular apparatus: Forms closed loop. [Transformational character] (0); Diverging laterally (1); Closed loop (2).
52. Perioral tentacular apparatus: Coiling direction. [Transformational character] (0); Anteriad (1); Posteriad (2).
53. Perioral tentacular apparatus: Adjustor muscle. Absent (0); Present (1).
54. Digestive tract: Prominent pharynx. Absent (0); Present (1).
55. Digestive tract: Radula. Absent (0); Present (1).
56. Digestive tract: Oesophageal folds. Absent (0); Present (1).
57. Digestive tract: Oral sphincter. Absent (0); Present (1).
58. Digestive tract: Foregut: Locomotory cilia. Absent (0); Present (1).
59. Digestive tract: Midgut: Subdivisions. Not subdivided (0); Functional subdivisions (1).
60. Digestive tract: Midgut: Glands. Absent (0); Present (1).
61. Digestive tract: Anus: Presence. Anus present: through-gut (0); Anus lost: digestive tract is blind sac (1).
62. Digestive tract: Anus: Location. [Transformational character] (0); Straight gut with [dorso] posterior anus (1); Anus migrated posteriad to create U-shaped gut (2); Anus opening to rear of pedal sole, causing slight U-shape to gut (3).
63. Digestive tract: Anus: Migration: Within ring of tentacles. [Transformational character] (0); Not within ring of tentacles (1); Anterior - within ring of feeding tentacles (2).
64. Digestive tract: Anus: Migration: Position. [Transformational character] (0); Left (1); Right (2); Dorsally (3); Ventrally (4).
65. Sclerites: Present in adult. Absent (0); Present (1).
66. Sclerites: Periodically shed and replaced. Absent (0); Present (1).
67. Sclerites: Bivalved. Scleritomous: without differentiated dorsal and ventral valves (0); Bivalved: scleritome dominated by prominent dorsal and ventral valve (1).
68. Sclerites: Accessory sclerites: Reduced. [Transformational character] (0); Accessory sclerites not reduced (1); Accessory sclerites absent: two valves only (2).
69. Sclerites: Accessory sclerites: Arrangement. Single field (0); Peripheral and medial fields with distinct sclerite arrangements (1).
70. Sclerites: Accessory sclerites: Symmetry. Dextral and sinistral forms absent (0); Occuring in dextral and sinistral forms (1).
71. Sclerites: Bivalved: Hinge line shape. [Transformational character] (0); Astrophic (1); Strophic (2).
72. Sclerites: Bivalved: Enclosing filtration chamber. No filtration chamber, or open chamber (0); Shells close to form enclosed filtration chamber (1).
73. Sclerites: Bivalved: Commissure: Exact correspondence of valve margins. [Transformational character] (0); Margins align exactly when valves closed (1); Margins of different shape or size (2).
74. Sclerites: Bivalved: Commissure: Sulcate. [Transformational character] (0); Rectimarginate (1); Uniplicate (2); Sulcate (3).
75. Sclerites: Bivalved: Commissure: Circular. [Transformational character] (0); Continuous round outline with no corners (save at the hinge);(1); Lateral margins linear (2).
76. Sclerites: Bivalved: Commissure: Lateral margins. [Transformational character] (0); Subparallel (1); Diverging (2); Initially diverging, becoming subparallel (3).
77. Sclerites: Bivalved: Apophyses. Absent (0); Present (1).
78. Sclerites: Bivalved: Apophyses: Morphology. [Transformational character] (0); Deltidiodont (1); Cyrtomatodont (2); Pseudodont (3).
79. Sclerites: Bivalved: Apophyses: Dental plates. Absent (0); Present (1).
80. Sclerites: Bivalved: Sockets. Absent (0); Present (1).
81. Sclerites: Bivalved: Socket ridges. Absent (0); Present (1).
82. Sclerites: Bivalved: Muscle scars: Ventral. Absent (0); Present (1).
83. Sclerites: Bivalved: Muscle scars: Ventral: Position. [Transformational character] (0); Posterolateral and medial attachments (1); Medial attachments only (2).
84. Sclerites: Bivalved: Muscle scars: Adjustor. Absent (0); Present (1).
85. Sclerites: Bivalved: Muscle scars: Dorsal adductors. [Transformational character] (0); Dispersed (1); Radially arranged (2); Quadripartite (3).
86. Sclerites: Bivalved: Muscle scars: Adductors: Position. [Transformational character] (0); Oblique (1); At high angle (2).
87. Sclerites: Bivalved: Muscle scars: Dermal muscles. Absent or weakly developed (0); Strongly developed (1).
88. Sclerites: Bivalved: Muscle scars: Unpaired median (levator ani). Absent (0); Present (1).
89. Sclerites: Bivalved: Muscle scars: Dorsal diductor. Absent (0); Present (1).
90. Sclerites: Bivalved: Muscle scars: Dorsal diductor: Position. [Transformational character] (0); Close to commissural plane (1); Oblique to commissural plane (2); At high angle to commissural plane (3).
91. Sclerites: Dorsal valve: Growth direction. [Transformational character] (0); Holoperipheral (1); Mixoperipheral (2); Hemiperipheral (3).
92. Sclerites: Dorsal valve: Posterior surface: Differentiated. Posterior face of dorsal valve not differentiated (0); Posterior face of dorsal valve forms distinct cardinal area or pseudointerarea (1).
93. Sclerites: Dorsal valve: Differentiated posterior surface: Morphology. Curved lateral profile (0); Planar lateral profile (1).
94. Sclerites: Dorsal valve: Posterior surface: Medial groove. Absent (0); Present (1).
95. Sclerites: Dorsal valve: Posterior surface: Notothyrium. Absent (0); Present (1).
96. Sclerites: Dorsal valve: Posterior surface: Notothyrium: Shape. [Transformational character] (0); Parallel-sided cleft (1); Triangular (2).
97. Sclerites: Dorsal valve: Posterior surface: Notothyrium: Chilidial plates. [Transformational character] (0); Open (1); Covered by chilidial plates (2).
98. Sclerites: Dorsal valve: Notothyrial platform. Absent (0); Present (1).
99. Sclerites: Dorsal valve: Medial septum. Absent (0); Present (1).
100. Sclerites: Dorsal valve: Cardinal shield. Absent (0); Present (1).
101. Sclerites: Dorsal valve: Cardinal processes. Absent (0); Present (1).
102. Sclerites: Dorsal valve: Cardinal teeth. Absent (0); Present (1).
103. Sclerites: Dorsal valve: Clavicles. Absent (0); Present (1).
104. Sclerites: Dorsal valve: Clavicles: Type of clavicles. [Transformational character] (0); Monoclavicle (1); Triclavicle (2).
105. Sclerites: Ventral valve: Growth direction. [Transformational character] (0); Holoperipheral (1); Mixoperipheral (2); Hemiperipheral (3).
106. Sclerites: Ventral valve: Relative size. [Transformational character] (0); Ventral valve markedly larger than dorsal valve (ventribiconvex) (1); Equivalve (subequally biconvex) (2); Dorsal valve markedly larger than ventral valve (dorsibiconvex) (3).
107. Sclerites: Ventral valve: Posterior surface: Differentiated. Posterior surface of shell not differentiated (0); Posterior surface of shell forms distinct cardinal area or pseudointerarea (1).
108. Sclerites: Ventral valve: Posterior margin growth direction. [Transformational character] (0); Inward-growing (1); Outward-growing (2).
109. Sclerites: Ventral valve: Posterior surface: Planar. Curved lateral profile (0); Planar lateral profile (1).
110. Sclerites: Ventral valve: Ligula. Absent (0); Present (1).
111. Sclerites: Ventral valve: Posterior surface: Extent. [Transformational character] (0); Low: Wider than deep (1); High: Deeper than wide (2).
112. Sclerites: Ventral valve: Posterior surface: Delthyrium. Absent (0); Present (1).
113. Sclerites: Ventral valve: Posterior surface: Delthyrium: Shape. [Transformational character] (0); Parallel sided (1); Triangular (2); Round (3).
114. Sclerites: Ventral valve: Posterior surface: Delthyrium: Shape: Aspect of rounded opening. [Transformational character] (0); Elongate: oval to rhombic (1); Essentially circular (2); Wider than long (3).
115. Sclerites: Ventral valve: Posterior surface: Delthyrium: Cover. [Transformational character] (0); Open (1); Covered, at least in part (2).
116. Sclerites: Ventral valve: Posterior surface: Delthyrium: Cover: Extent. [Transformational character] (0); Covered only partially; partially open (1); Completely covered (2).
117. Sclerites: Ventral valve: Posterior surface: Delthyrium: Cover: Identity. [Transformational character] (0); Pseudodeltidium (1); Deltidial plate(s) (2); Continuation of shell (3).
118. Sclerites: Ventral valve: Posterior surface: Delthyrium: Pseudodeltidium: Shape. [Transformational character] (0); Concave (1); Convex (2).
119. Sclerites: Ventral valve: Posterior surface: Delthyrium: Pseudodeltidium: Hinge furrows. Absent (0); Present (1).
120. Sclerites: Ventral valve: Umbonal perforation. Umbo imperforate (or ventral valve absent) (0); Umbonal perforation (1).
121. Sclerites: Ventral valve: Umbonal perforation: Shape. [Transformational character] (0); Circular (or subcircular) (1); Rhombic to oval (2); Arising through decollation (3).
122. Sclerites: Ventral valve: Colleplax, cicatrix or pedicle sheath. Absent (0); Present (1).
123. Sclerites: Ventral valve: Median septum. Absent (0); Present (1).
124. Sclerites: Ornament: Concentric ornament. [Transformational character] (0); Smooth, or growth lines only (1); Concentric ornament present (2).
125. Sclerites: Ornament: Concentric ornament: Symmetry. [Transformational character] (0); Asymmetric fila, with outer faces (1); Symmetric fila (2).
126. Sclerites: Ornament: Radial ornament. Absent (0); Present (1).
127. Sclerites: Ornament: Shell-penetrating spines. Absent (0); Present (1).
128. Sclerites: Composition: Mineralogy. [Transformational character] (0); Organic (non-mineralized) (1); Phosphatic (2); Calcitic (3); Aragonitic (4).
129. Sclerites: Composition: Cuticle or organic matrix. [Transformational character] (0); GAGs, chitin and collagen (1); Glycoprotein (2).
130. Sclerites: Composition: Incorporation of sedimentary particles. Absent (0); Present (1).
131. Sclerites: Composition: Periostracum: Flexibility. [Transformational character] (0); Flexible (1); Inflexible (2).
132. Sclerites: Composition: Microstructure: Number of distinct layers. [Transformational character] (0); Single microstructural layer (1); Two microstructurally differentiated layers (2); Inner and outer laminae enclosing medial void (3); Three microstrurally differentiated layers (4).
133. Sclerites: Composition: Microstructure: Format. [Transformational character] (0); Laminated (1); Fibrous bundles (2); Nacreous / crossed lamellar (3).
134. Sclerites: Structure: Stratiform lamellae expressed at surface. [Transformational character] (0); Lamellae not expressed at surface (1); Lamellae correspond to external shell ornament (2).
135. Sclerites: Structure: Stratiform laminae separated. [Transformational character] (0); Contiguous stratified layer (1); Laminae separated by organic layers or voids (2).
136. Sclerites: Structure: Stratiform laminae with polygonal ornament. [Transformational character] (0); Absent (1); Present (2).
137. Sclerites: Structure: Canals. Absent (0); Present (1).
138. Sclerites: Structure: Punctae. Absent (0); Present (1).
139. Sclerites: Structure: Pseudopunctae. Absent (0); Present (1).
140. Sclerites: Structure: External polygonal ornament. Absent (0); Present (1).
141. Gametes: Gonocoel. Absent (0); Retroperineal gonads (1).
142. Gametes: Ovary wall saccular. Plain (0); Saccular (1).
143. Gametes: Testis wall saccular. Plain (0); Saccular (1).
144. Gametes: Asexual reproduction. Never exhibited (0); Frequently exhibited (1).
145. Gametes: Sexes. [Transformational character] (0); Gonochoristic (1); Hermaphroditic (2).
146. Gametes: Fertilization. [Transformational character] (0); External (1); Internal (2).
147. Gametes: Egg: Size. [Transformational character] (0); Small: < 100 um, little yolk (1); Large: > 110 um, much yolk (2).
148. Gametes: Egg: Protective membrane. Absent (0); Present (1).
149. Gametes: Egg: Site of maturation. [Transformational character] (0); Body cavity (1); Mantle canals (2); Ovicell (3).
150. Gametes: Spermatozoa: Nucleus: Shape. Equant: length comparable to width (0); Elongate: length exaggerated relative to width (1).
151. Gametes: Spermatozoa: Anterior nuclear fossa. Absent (0); Present (1).
152. Gametes: Spermatozoa: Acrosome: Shape. [Transformational character] (0); Pear-shaped (1); Needle-shaped (2); Disc-shaped (3); Conical (4).
153. Gametes: Spermatozoa: Acrosome: Differentiated internally. No internal differentiation (0); Acrosome differentiated internally (1).
154. Gametes: Spermatozoa: Acrosome: Sub-acrosomal space. Absent (0); Present (1).
155. Gametes: Spermatozoa: Mid-piece. Multiple mitochondria (0); Single ring-shaped mitochondrion (1).
156. Gametes: Spermatozoa: Centrioles: Orientation. Orthogonal (0); Parallel (1).
157. Gametes: Spermatozoa: Centrioles: Fusion. Discrete (0); Fused (1).
158. Gametes: Spermatozoa: Satellite fibre complex. Annulus not associated with satellite fibres (0); Annulus associated with satellite fibres (1).
159. Gametes: Spermatozoa: Mitochondria: Shape. [Transformational character] (0); Spherical to subspherical (1); Rods (2); Elongate, sac-like (3).
160. Gametes: Spermatozoa: Mitochondria: Cristae: Configuration. Unmodified (0); Arranged in parallel plates (1).
161. Gametes: Spermatozoa: Mitochondria: Midpiece. [Transformational character] (0); Extremely short (1); Long (2); Forms continuous sheath (3).
162. Embryo: Micromere size. [Transformational character] (0); Similar to macomeres (1); Small relative to macromeres (2).
163. Embryo: Cleavage: Equal. [Transformational character] (0); Unequal (1); Equal (2).
164. Embryo: Cleavage: Cross pattern. Absent (0); Cross, whether “molluscan” or “annelid” (1).
165. Embryo: Cleavage: Polar lobe formation. [Transformational character] (0); Absent (1); Present (2).
166. Embryo: Cleavage: Spiral. [Transformational character] (0); Absent (1); Present (2).
167. Embryo: Origin of mesoderm. [Transformational character] (0);4d cell, from the blastopore ridge, or as ectomesoderm (1); Archenteron (2).
168. Larva: Apical organ: Muscles extending to the hyposphere. Absent (0); Present (1).
169. Larva: Apical organ: Serotonergic cells. [Transformational character] (0); Two flask-shaped cells (1); Four flask-shaped cells (2); Cluster of c. eight flask-shaped cells (3); Aggregation of multiple cells of multiple types (4).
170. Larva: Apical organ: Develops into adult brain. Brain has other origin (0); Adult brain derived from larval apical organ / apical pole (1); [Transformational character] (2).
171. Larva: Brain persists into adulthood. [Transformational character] (0); Brain lost (1); Brain retained to adulthood (2).
172. Larva: Origin of body cavity. [Transformational character] (0); Mesenchyme (1); Coelom (2).
173. Larva: Formation of coelomoducts. [Transformational character] (0); Outgrowth (1); Ingrowth (2).
174. Larva: Foot. Absent (0); Present (1).
175. Larva: Foot: Pedal gland. Absent (0); Present (1).
176. Larva: Coelom: Paired. Absent (0); Paired coelom originating from two teloblasts derived from 4d (1).
177. Larva: Coelom: Paried: Includes pericardium. Paired coelom absent, or does not include pericardium (0); Paired coelom includes pericardium (1).
178. Larva: Feeding. [Transformational character] (0); Lecithotrophic (or otherwise non-feeding) (1); Planktotrophic (or otherwise feeding) (2).
179. Larva: Cilia: Metatroch. Absent (0); Present (1).
180. Larva: Cilia: Telotroch. Absent (0); Present (1).
181. Larva: Cilia: Ciliated food groove. Absent (0); Present (1).
182. Larva: Cilia: Ciliary bands: Downstream. Absent (0); Present (1).
183. Larva: Cilia: Ciliary bands: Upstream. Absent (0); Present (1).
184. Larva: Cilia: Adoral ciliary band. Absent (0); Present (1).
185. Larva: Cilia: Nerve ring underlying ciliated larval swimming organ. Absent (0); Present (1).
186. Ciliary ultrastructure: Accessory centriole. Absent (0); Present (1).
187. Ciliary ultrastructure: Aggregation of granules below basal plate. Absent (0); Present (1).
188. Ciliary ultrastructure: Basal foot: Radiating tubular fibres. Absent (0); Present (1).
189. Ciliary ultrastructure: Basal plate. [Transformational character] (0); Thin (1); Blurry (2).
190. Ciliary ultrastructure: Brushborder of microvilli. Absent (0); Present (1).
191. Ciliary ultrastructure: Centriolar triplet derivative in basal body. [Transformational character] (0); 9 + 2 pattern (1); 9 + 3 pattern (2).
192. Ciliary ultrastructure: Ciliary necklace with connecting strands. Absent (0); Present (1).
193. Ciliary ultrastructure: Compound cilia: Presence. Absent (0); Present (1).
194. Ciliary ultrastructure: Compound cilia: Origin. [Transformational character] (0); Several monociliate cells (1); On multiciliated cell (2).
195. Ciliary ultrastructure: Glycocalyx ultrastructure. [Transformational character] (0); Homogeneous (1); Layered (2).
196. Ciliary ultrastructure: Microvilli on epidermal surface: Branched. Unbranched (0); Branched (1).
197. Ciliary ultrastructure: Vertical ciliary rootlet: Length. [Transformational character] (0); Short (1); Long (2).
198. Ciliary ultrastructure: Vertical ciliary rootlet: Shape. [Transformational character] (0); Conical (1); Flat (2).
199. Ciliary ultrastructure: Secondary ciliary rootlet: Presence. Absent (0); Present (1).
200. Ciliary ultrastructure: Secondary ciliary rootlet: Length. [Transformational character] (0); Short (1); Long (2).
201. Ciliary ultrastructure: Secondary ciliary rootlet: Shape. [Transformational character] (0); Conical (1); Flat (2).
202. Nephridia: Podocytes. Absent (0); Present (1).
203. Nephridia: Rhogocytes. Absent (0); Present (1).
204. Nephridia: Serve as excretory organs. No (0); Yes (1).
205. Nephridia: Protonephridia. Absent (0); Present (1).
206. Nephridia: Metanephridia. Absent (0); Present (1).
207. Cuticle: Layers. Simple (i.e. glycocalyx) (0); Distinct epicuticle and endocuticle (1).
208. Cuticle: Composition. [Transformational character] (0); Chitinous (1); Collagen (2).
209. Cuticle: Fibrous layer with thick fibrils. Absent (0); Present (1).
210. Cuticle: Homogeneous layer. Absent (0); Present (1).
211. Cuticle: Resilience. Labile (0); Robust (1); [Transformational character] (2).
212. Cuticle: Microvilli. Absent (0); Microvilli present in the cuticle (1).
213. Muscles: Cytology. [Transformational character] (0); Smooth (1); Obliquely striated (2); Smooth on abfrontal face; striated on frontal face (3).
214. Muscles: Histology. Fibre-type (0); Epithelially organized (1); [Transformational character] (2).
215. Glands: Pedal gland. Absent (0); Present (1).
216. Glands: Paired pharyngeal diverticulae. Absent (0); Present (1).
217. Nervous system: Orthogonal. Not orthogonal (0); Orthogonal (1).
218. Nervous system: Glial system. Absent (0); Present (1).
219. Nervous system: Buccal nerve ring. Absent (0); Present (1).
220. Nervous system: Anterior nerve loop. Absent (0); Present (1).
221. Nervous system: Formation of ganglia. [Transformational character] (0); From cerebral region (1); In situ (2); Invagination of epithelium (3).
222. Nervous system: Cerebral ganglia: Presence. Absent (0); Present (1).
223. Nervous system: Cerebral gangila: Fused. [Transformational character] (0); Pair of distinct ganglia (1); Single ganglion, or fused ganglia (2).
224. Nervous system: Nerve cords. [Transformational character] (0); Ventral nerve cords only (1); Tetraneury: one pair of ventral and one pair of lateral nerve cords (2).
225. Nervous system: Ventral longitudinal nerves. Separate (0); Paired or secondarily fused (1).

#### New characters

226. Body: morphology. Non-vermiform (0); Vermiform (1). [[Non-vermiform in crown-group brachiopods]]
227. Trunk: shape. Tubular (0); Conical (1). [[Conical in *Conicula*, *Yuganotheca*, and hyoliths]]
228. Trunk region: biomineralization. Absent (0); Present (1). [[Present in hyoliths, *Yuganotheca*, and crown-group brachiopods]]
229. Median collar or ring connecting trunk and lophophoral regions: Presence. Absent (0); Present (1). [[Present in *Conicula* and *Yuganotheca*]]
230. Lophophoral region: biomineralization. Absent (0); Present (1). [[Present in hyoliths and crown-group brachiopods]]
231. Mantle surrounding a lophophoral chamber: Presence. Absent (0); Present (1). [[Present in *Conicula*, brachiopods, and associated brachiopod stem groups]]
232. Mantle: morphology. Undivided (0); Bilobed (1). [[Undivided mantle in *Conicula*, bilobed mantle in *Yuganotheca* and crown-group brachiopods]]

#### Alternative evolutionary affinities

Recently, Zhao *et al*. (2021) proposed a stem-group medusozoan cnidarian affinity of *C*. *striata* in a preprint. However, this phylogenetic position is strongly contradicted to the presence of a through and U-shaped gut in *C*. *striata*, which is clearly revealed by the fossil material represented here (Fig. 3l–n). Although the stomach was preserved in some of the specimens figured in Zhao *et al*. (2021), unfortunately it was interpreted incorrectly as “peduncle chamber”. As a through and U-shaped gut is hardly a feature of any known cnidarian-grade metazoans, the cnidarian affinity of *C*. *striata* inferred by Zhao *et al*. (2021) is highly questionable. Under this working model, their interpretations of other anatomical structures shall be taken into careful reconsideration.

---

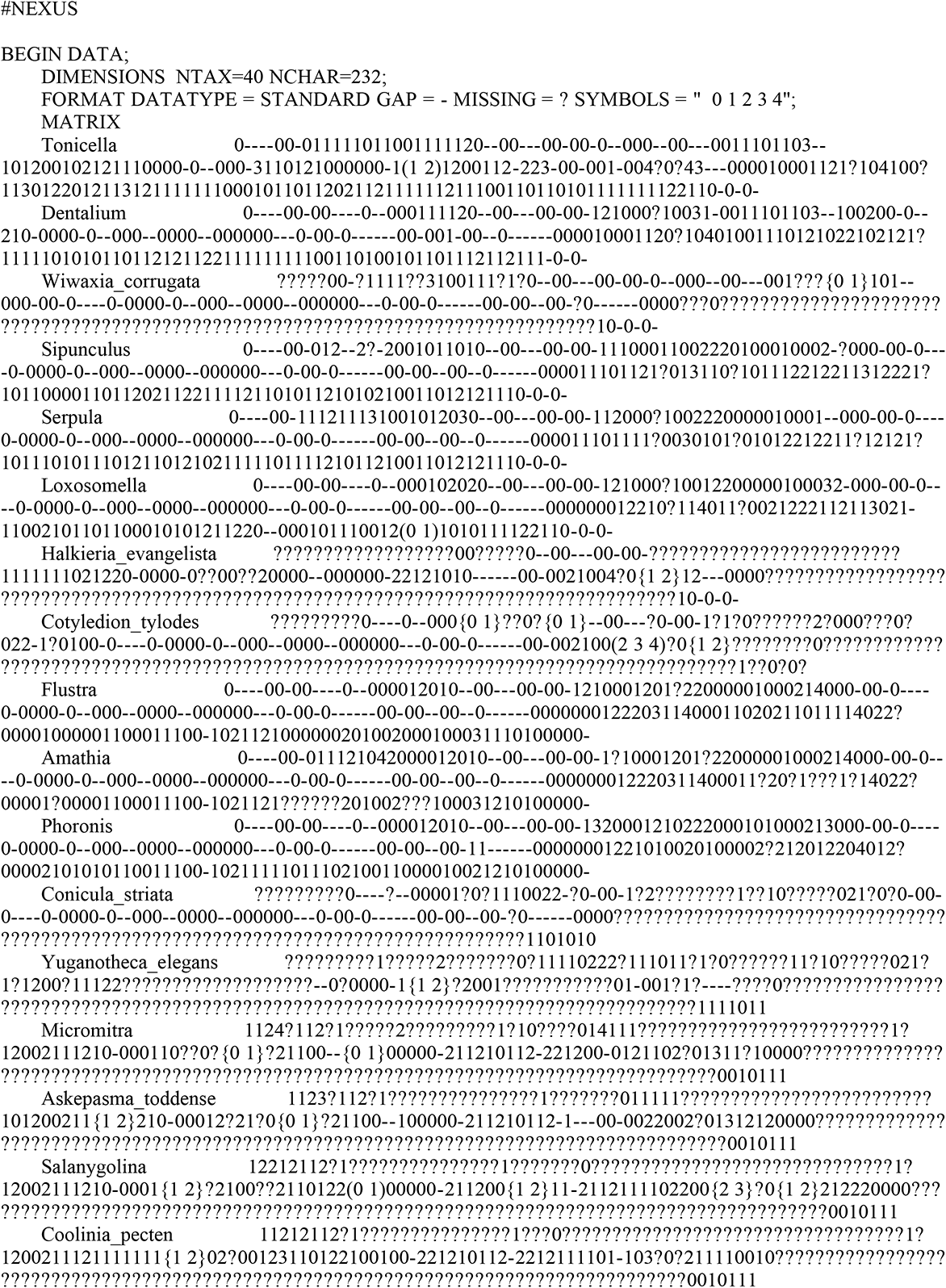
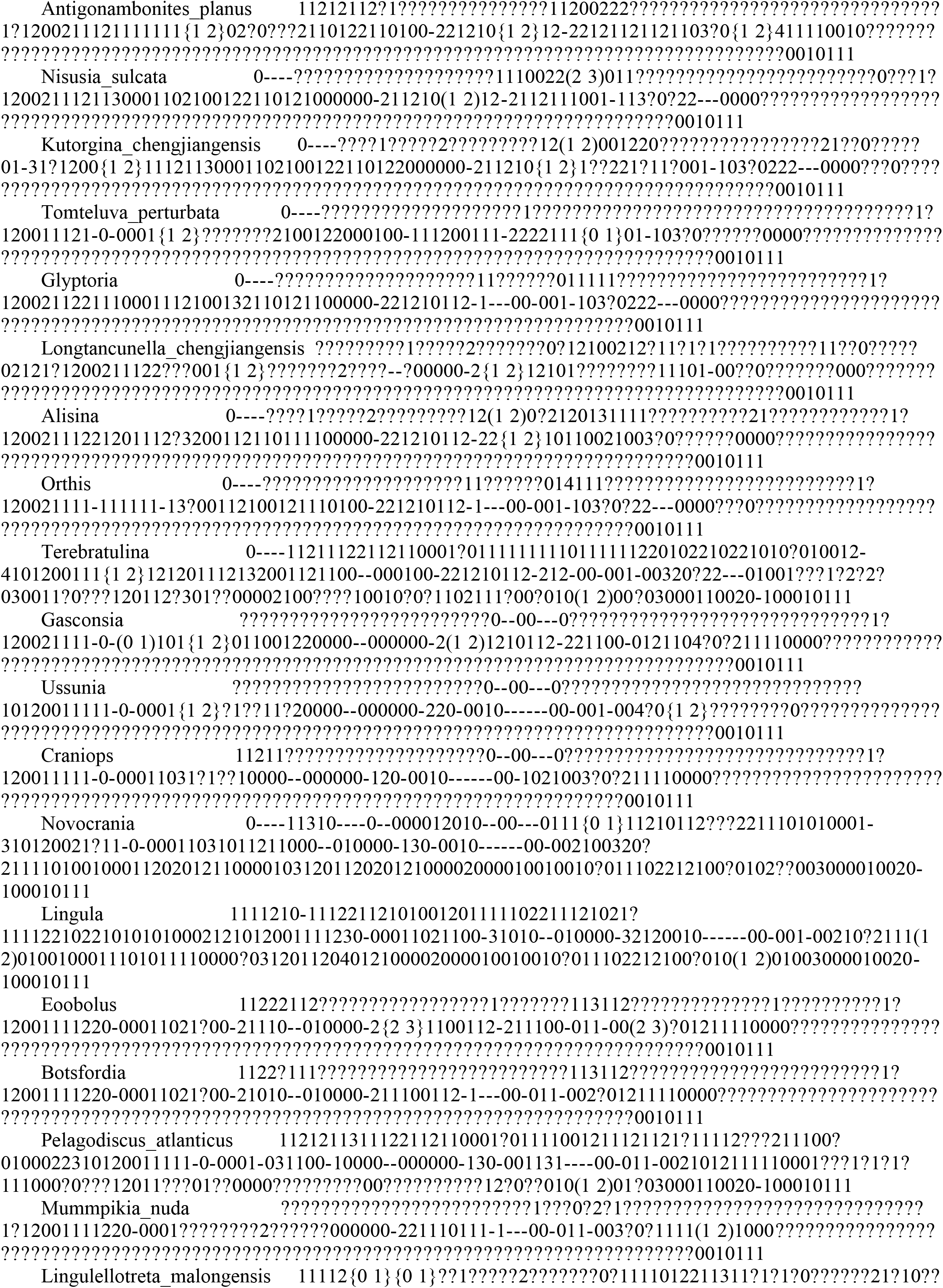
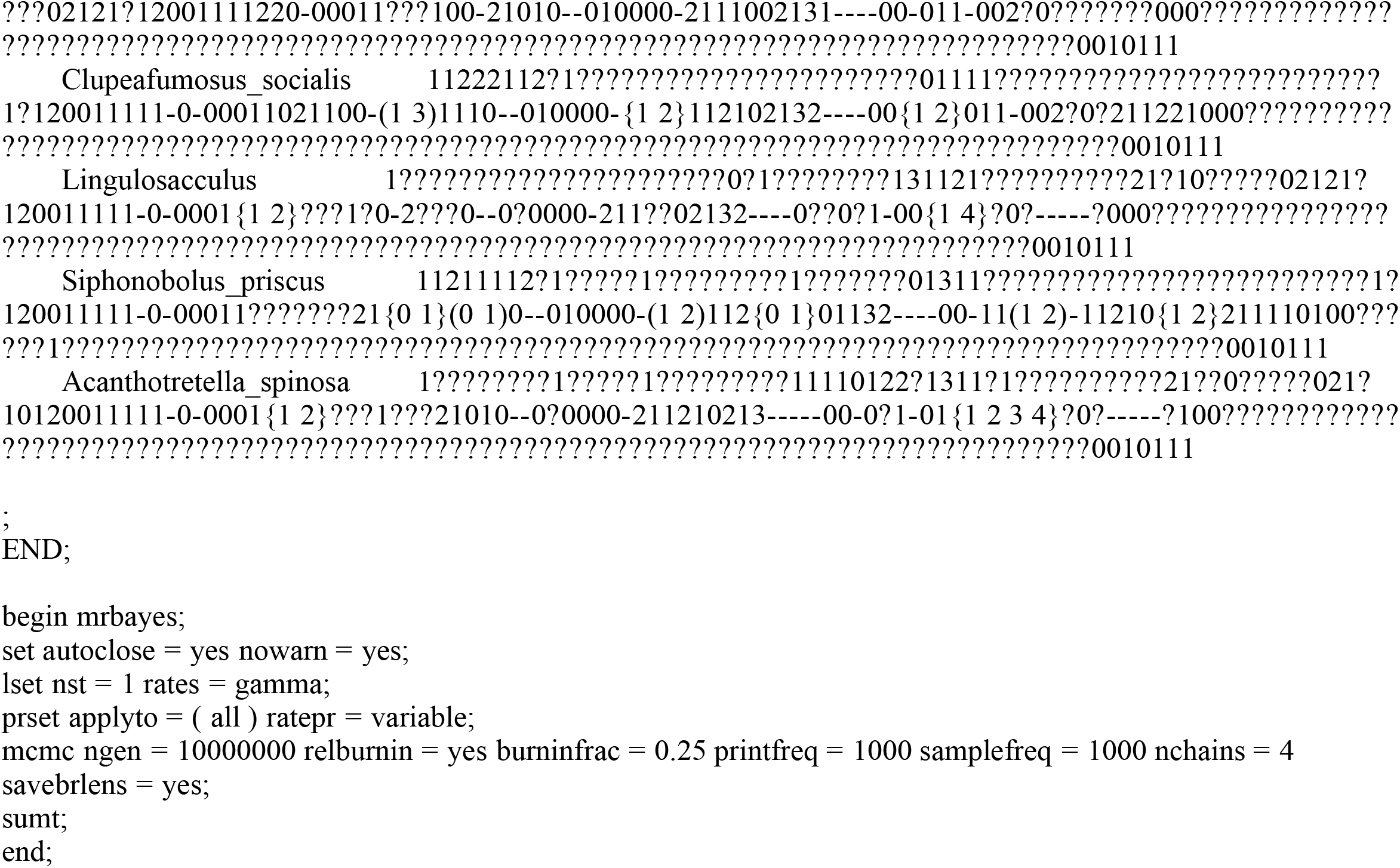

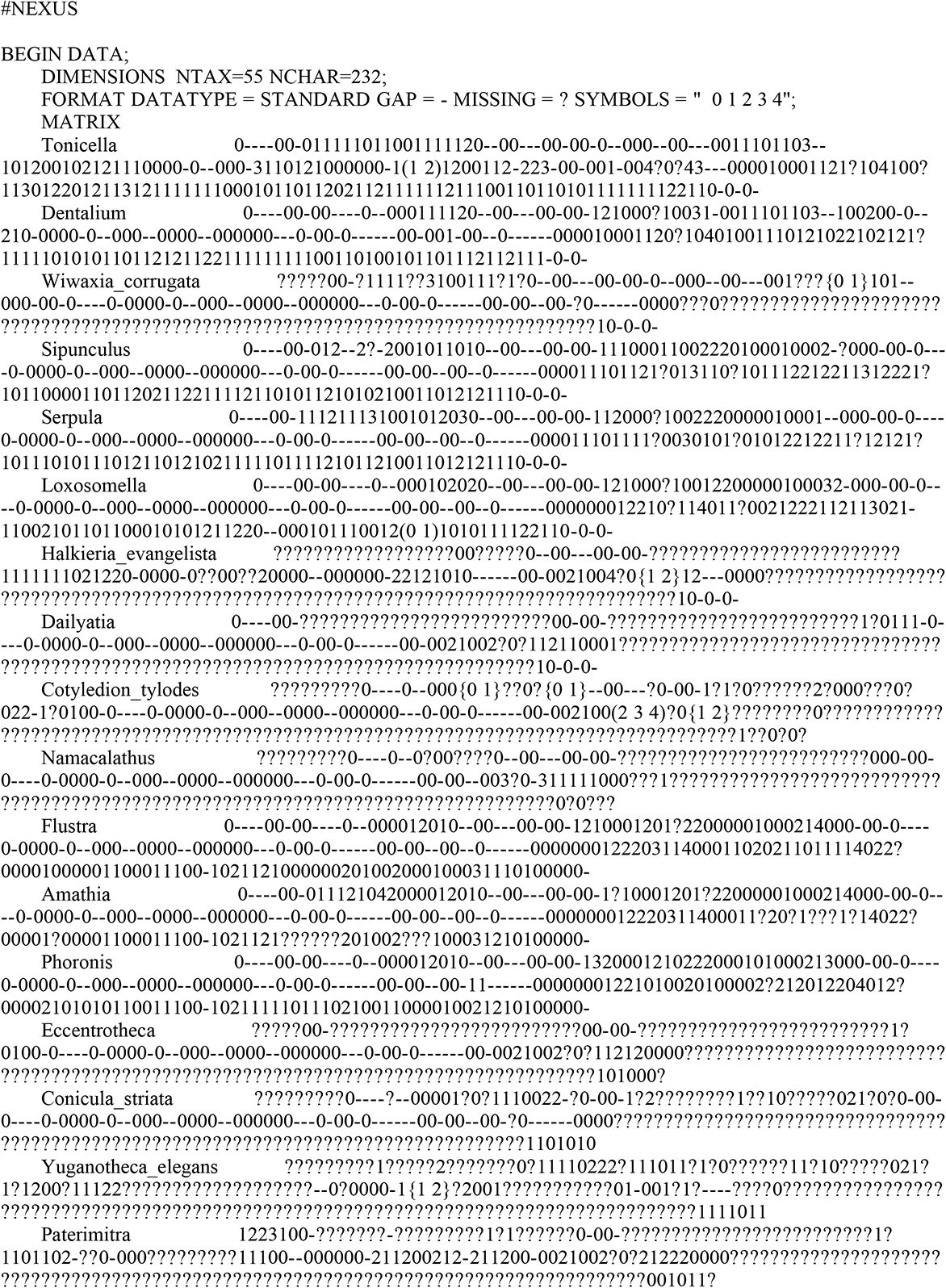
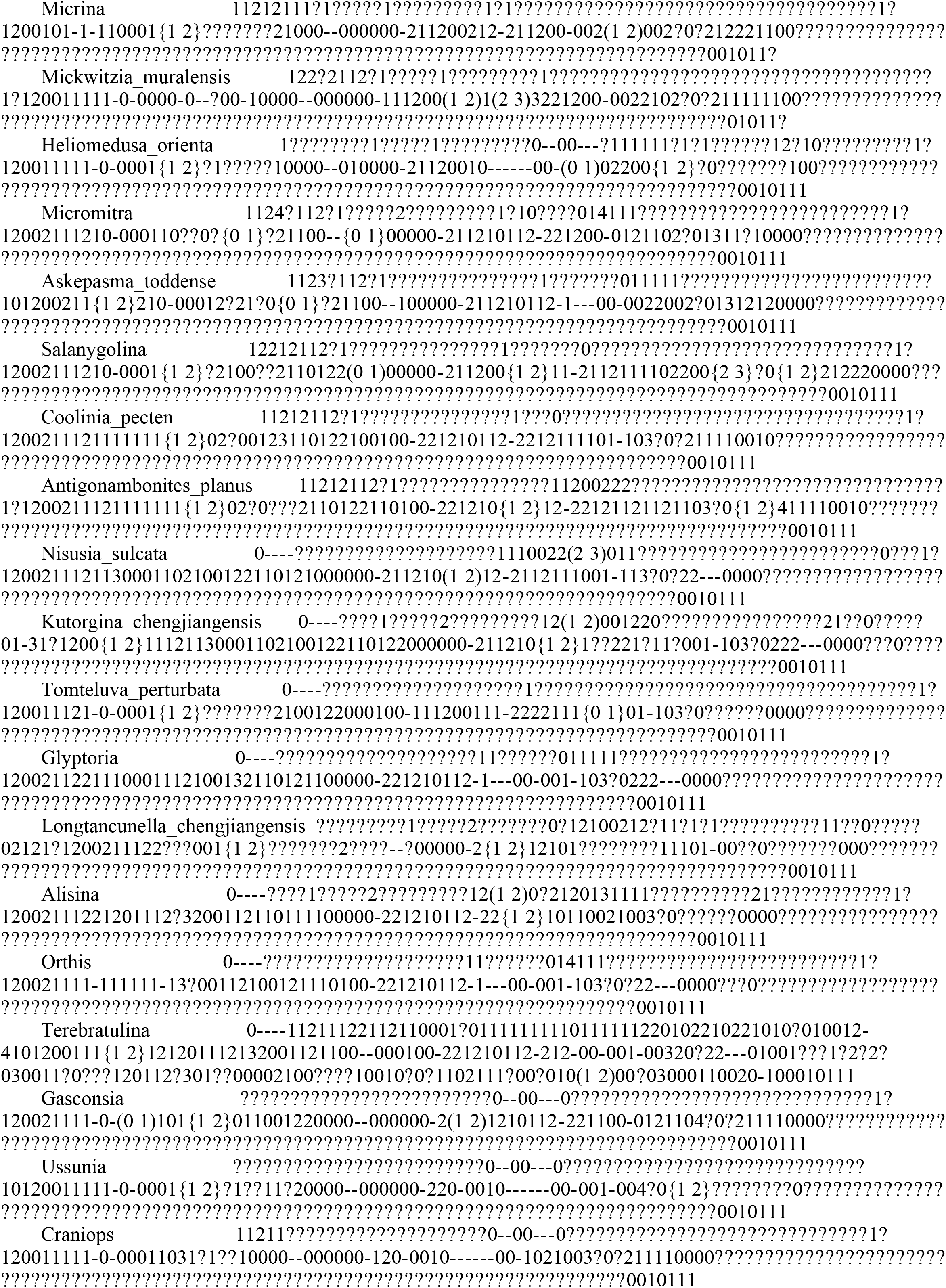
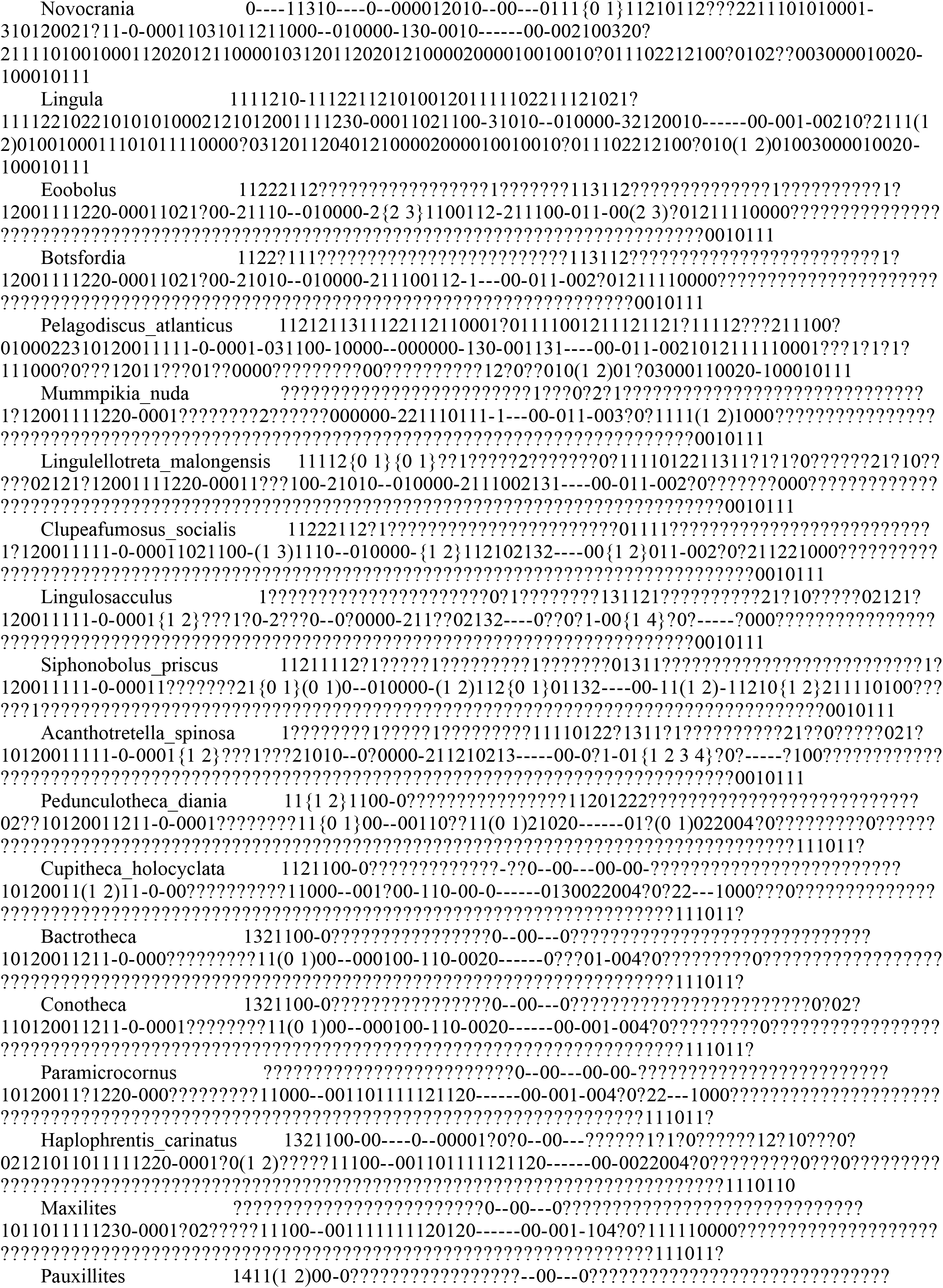
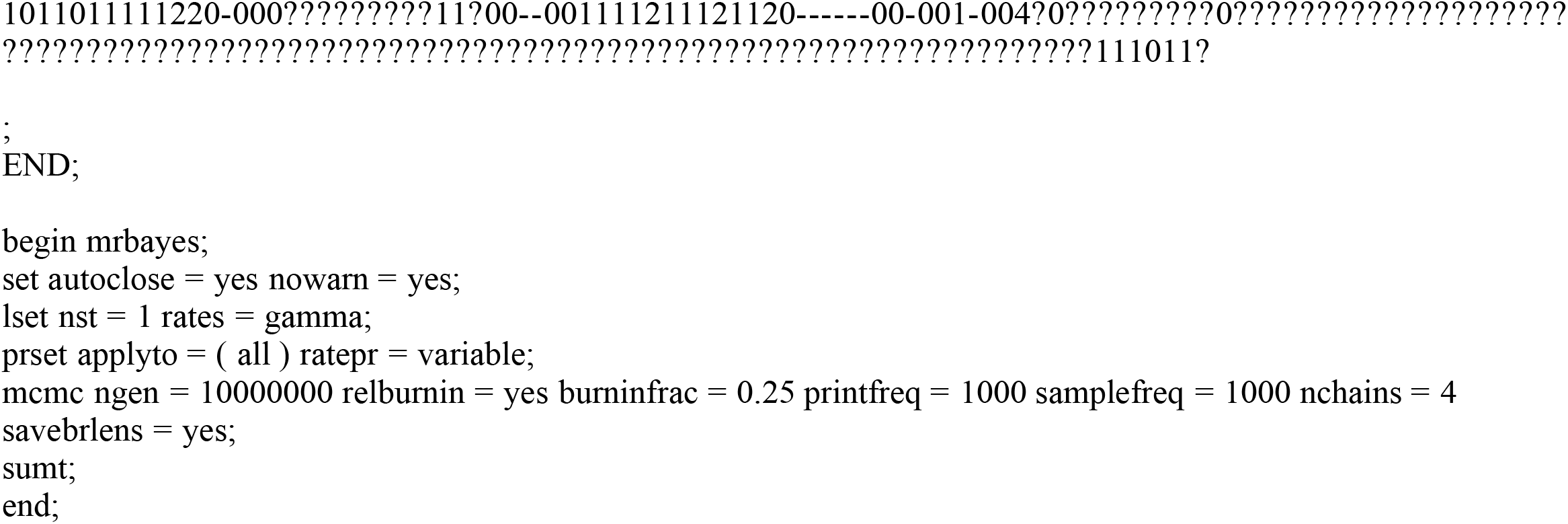

## Notes

### Competing Interest Statement

The authors have declared no competing interest.

